# Dual carbon sequestration with photosynthetic living materials

**DOI:** 10.1101/2023.12.22.572991

**Authors:** Dalia Dranseike, Yifan Cui, Andrea S. Ling, Felix Donat, Stéphane Bernhard, Margherita Bernero, Akhil Areeckal, Xiao-Hua Qin, John S. Oakey, Benjamin Dillenburger, André R. Studart, Mark W. Tibbitt

**Affiliations:** Macromolecular Engineering Laboratory, Department of Mechanical and Process Engineering, ETH Zurich, Zurich, CH; Digital Building Technologies, Department of Architecture, ETH Zurich, Zurich, CH; Laboratory of Energy Science and Engineering, Department of Mechanical and Process Engineering, ETH Zurich, Zurich, CH; Institute for Biomechanics, Department of Health Sciences and Technology, ETH Zurich, Zurich, CH; Department of Chemical and Biomedical Engineering, University of Wyoming, Laramie, Wyoming, US; Complex Materials, Department of Materials, ETH Zurich, Zurich, CH

**Keywords:** living materials, CO_2_ sequestration, 3D printing

## Abstract

Natural ecosystems offer efficient pathways for carbon sequestration, serving as a resilient approach to remove CO_2_ from the atmosphere with minimal environmental impact. However, the control of living systems outside of their native environments is often challenging. Here, we engineered a photosynthetic living material for dual CO_2_ sequestration by immobilizing photosynthetic microorganisms within a printable polymeric network. The carbon concentrating mechanism of the cyanobacteria enabled accumulation of CO_2_ within the cell, resulting in biomass production. Additionally, the metabolic production of OH^-^ ions in the surrounding medium created an environment for the formation of insoluble carbonates via microbially-induced calcium carbonate precipitation (MICP). Digital design and fabrication of the living material ensured sufficient access to light and nutrient transport of the encapsulated cyanobacteria, which were essential for long-term viability (more than one year) as well as efficient photosynthesis and carbon sequestration. The photosynthetic living materials sequestered approximately 2.5 mg of CO_2_ per gram of hydrogel material over 30 days via dual carbon sequestration, with 2.2 ± 0.9 mg stored as insoluble carbonates. Over an extended incubation period of 400 days, the living materials sequestered 26 ± 7 mg of CO_2_ per gram of hydrogel material in the form of stable minerals. These findings highlight the potential of photosynthetic living materials for scalable carbon sequestration, carbon-neutral infrastructure, and green building materials. The simplicity of maintenance, coupled with its scalability nature, suggests broad applications of photosynthetic living materials as a complementary strategy to mitigate CO_2_ emissions.

## Introduction

Biological ecosystems, such as forests, aquatic systems, and wetlands, offer efficient pathways for carbon sequestration (storing carbon in a carbon pool) and conversion into carbon-based materials^1^. Natural systems operate under ambient conditions with sunlight and commonly available small molecules as their sole inputs. Further, living systems can sense, self-repair, and respond to their surroundings, making them resilient to environmental changes^1,2^. Biological carbon sequestration, for example via afforestation or the growth of marine phytoplankton and algae, is also cost-efficient and environmentally-friendly^3^. In this context, natural carbon sequestration can serve as a passive, low-impact complement to industrial carbon sequestration, which normally requires specific, extreme, and energy-intensive conditions^1,4^ and proximity to large emission sources^5^. However, natural carbon sequestration is typically slower than industrial carbon sequestration, and the control of living systems outside of their native environments is often challenging^5,6^.

Strategies to engineer living systems for active CO_2_ sequestration would provide an additional approach to mitigate the accumulation of human-generated CO_2_ in the atmosphere. The CO_2_ concentrating mechanism of many photosynthetic microorganisms accumulates CO_2_ within the cell body up to 1000-fold above ambient levels^6,7^. Subsequently, concentrated carbon can be fixed in the form of biomass generated during growth^8,9^. In addition to biomass production, microbially-induced calcium carbonate precipitation (MICP) in certain species can sequester CO_2_ irreversibly in the form of inorganic carbonate precipitates. MICP proceeds via multiple metabolic pathways, including ureolysis, sulfate reduction, and denitrification^10,11^. In some organisms, MICP can occur as a direct by-product of photosynthesis, whereby the inorganic precipitates effectively act as an additional carbon sink^12^, enabling dual carbon sequestration. In this context, immobilizing photosynthetic microorganisms, such as algae and cyanobacteria, within a support matrix may provide an approach to drive biological CO_2_ sequestration in the form of engineered photosynthetic living materials via dual carbon sequestration.

To date, engineered living materials have primarily been used for applications in biomedicine, sustainable materials production, and as living building materials^13–17^. For example, MICP has been exploited, primarily via ureolysis, to mechanically reinforce living materials based on the in situ formation of a stiff mineral phase^18^. Robust composites were produced via biomineralization using ureolytic MICP within a cellulose matrix^19,20^. Similarly, precipitates deposited in porous materials filled cracks and improved mechanical properties of composite building structures as well as consolidated soils^21^. Ureolytic MICP is attractive due to its short incubation period (typically 1-4 days), resistance to contamination, and rapid biomineralization; however, it poses substantial environmental concerns due to the associated production of large amounts (1-2 equimolar) of ammonia^22^. Further, ureolytic MICP requires a constant supply of urea and only proceeds in a narrow range of environmental conditions^23–25^. These challenges restrict the use of ureolytic MICP for long-term CO_2_ sequestration^26^. Many of these limitations can be addressed with photosynthetic MICP, which requires no additional feedstocks and produces no toxic by-products^18,25^. Recently, photosynthetic MICP was used to design living building materials that mineralized over time^17^. While photosynthetic living materials have been explored for carbon sequestration via reversible biomass accumulation^9^, they have not been explored for CO_2_ sequestration via biomass accumulation *and* irreversible MICP using atmospheric CO_2_ as the main carbon source and light as the sole source of energy.

Drawing inspiration from natural systems, we engineered photosynthetic living materials for dual CO_2_ sequestration by immobilizing MICP-capable photosynthetic microorganisms within a printable polymeric network. Cyanobacterium *Synechococcus* sp. strain PCC 7002 (herein referred to as PCC 7002) was chosen for this work as it possesses many prominent advantages. This strain can synthesize complex carbohydrates using light, inorganic nutrients found in seawater, and atmospheric CO_2_ as the main carbon source. Further, PCC 7002 is capable of photosynthetic MICP, exhibits a fast doubling time (∼2.6 h under optimal conditions), and tolerates variations in light intensity and osmotic pressure^27–29^. A bio-inert Pluronic F-127 (F127)-based hydrogel was used as the support matrix, which was processed via additive manufacturing into engineered 3D living structures^30^. The transparency of the hydrogel facilitated light penetration and cell growth within the structure. Inspired by aquatic structures and capillary-driven fluid flow, photosynthetic living materials were designed as fine-scale open lattices, branched forms, and discrete pillars to enhance access to light and nutrient exchange and to augment photosynthetic biomass generation^31,32^. Dual carbon sequestration via biomass generation and insoluble carbonate formation proceeded over the lifecycle (beyond one year) of the bio-printed structures. The mineral phase mechanically reinforced the living materials and stored sequestered carbon in a more stable form. This work provides a strategy to engineer photosynthetic living materials as a scalable CO_2_ sequestration method to complement ongoing strategies to mitigate atmospheric CO_2_ accumulation.

## Results

### Engineering living materials for dual carbon sequestration

We engineered photosynthetic living materials for dual carbon sequestration by exploiting the carbon concentrating mechanism of cyanobacteria strain PCC 7002 **(Figure 1a)**. Dual CO_2_ sequestration proceeds via both reversible biomass accumulation and irreversible mineral precipitation. For carbon concentration to occur, CO_2_ from the surrounding environment dissolves in aqueous solutions and forms bicarbonate ions (HCO_3_^-^) that are then transported to the bacterial carboxysome^33^. Here, carbonic anhydrase (CA) within the carboxysome converts the bicarbonate species into a hydroxyl ion (OH^-^) and CO_2_. The OH^-^ is excreted to the exterior of the bacterium and the CO_2_ is fixed by ribulose-1,5-bisphosphate carboxylase/oxygenase (RuBisCo) during photosynthesis into two molecules of 3-phosphoglycerate, which is enzymatically converted into sugars that support cell growth and biomass production. Due to the metabolic production of OH^-^, the local pH value of the local medium surrounding the bacteria increases. This increase in pH, combined with negatively charged extracellular polysaccharides on the bacterial membrane, creates a favorable environment for the nucleation and formation of insoluble carbonates^31^. In the presence of divalent cations, such as Ca^2+^ and Mg^2+^, CO_3_^2-^ in the medium is consumed and fixed irreversibly into calcium or magnesium carbonates. Chemical equilibrium then favors the dissolution of additional atmospheric CO_2_ into the culture medium, continuously driving the dual carbon sink.

**Figure 1.**
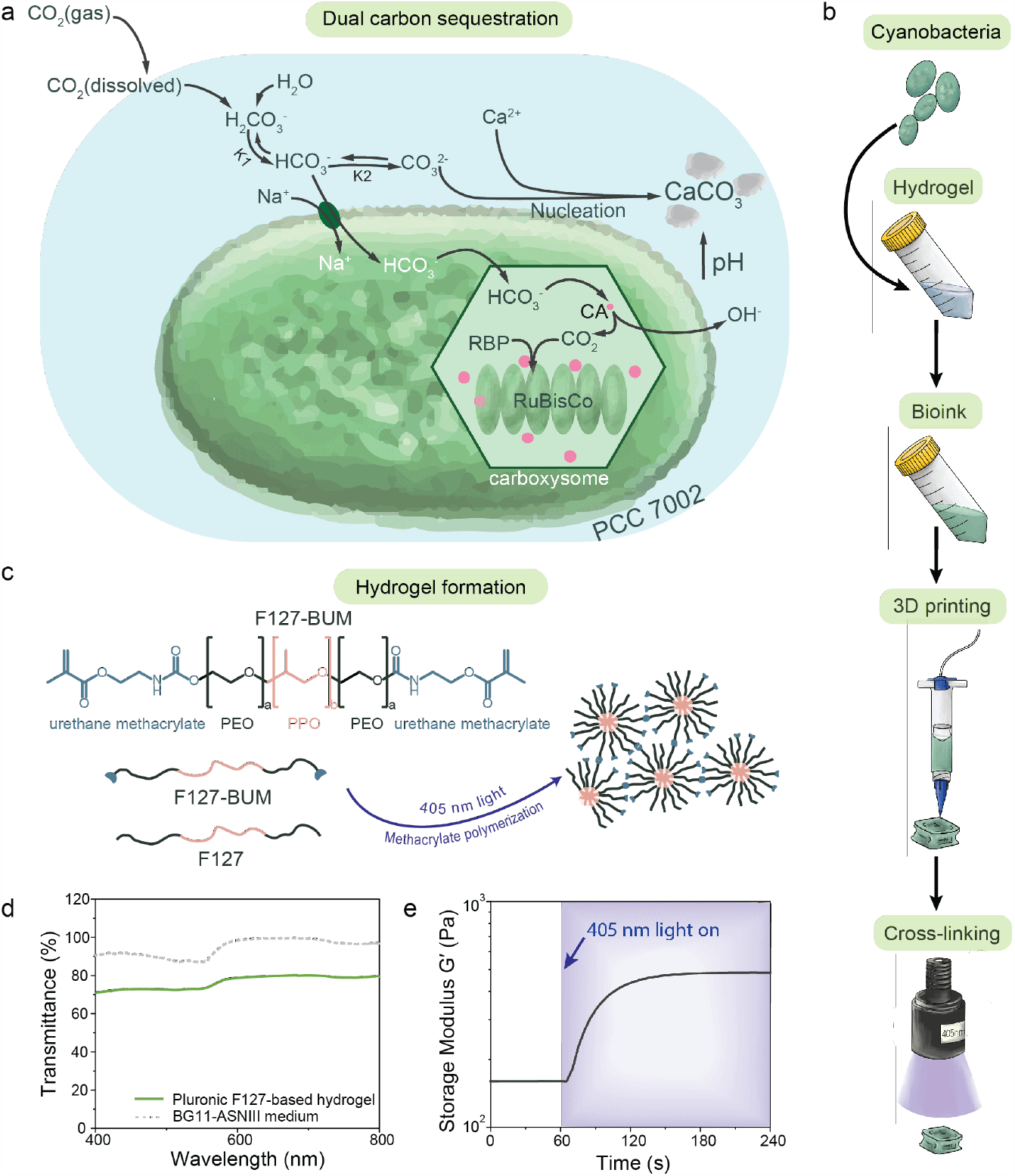
Preparation of photosynthetic living materials for dual carbon sequestration. **a)** PCC 7002 are capable of CO2 sequestration via biomass accumulation and microbially induced carbonate precipitation (MICP) during photosynthesis using dissolved CO_2_ as the major carbon source. **b)** Photosynthetic living materials were fabricated by combining PCC 7002 with a hydrogel bioink, that was structured via 3D printing and stabilized via photopolymerization. **c)** The support hydrogel was composed of a blend of F127 and F127-BUM, which was photo-cross-linked to form the final stable constructs. **d)** The hydrogel permitted visible light penetration to support the photosynthesis of encapsulated PCC 7002. **e)** The storage modulus (G′) increased upon photo-cross-linking under 405 nm (I = 8 mW cm^-1^) light rendering the stable final materials.

Cyanobacteria PCC 7002 was encapsulated in a hydrogel matrix to fabricate photosynthetic living materials capable of dual carbon sequestration **(Figure 1b)**. A synthetic polymeric hydrogel based upon Pluronic F127 (F127) was chosen for the encapsulation of cyanobacteria due to its bioinert nature and its processing versatility. F127-based hydrogels have been used in the design of engineered living materials due to the facile diffusion of most small molecules through it^34^. Functionalized F127-bis urethane methacrylate (F127-BUM) can be photo-cross-linked either post-printing or processed directly via light-based additive manufacturing for long-term structural stability **(Figure 1c)**. In order to structure the photosynthetic living materials using digital fabrication, we designed a bioink (13.2 wt% F127 and 7.3 wt% F127-BUM) that maintained high viability of encapsulated PCC 7002 and printability via direct ink writing and light-based additive manufacturing. Rationally designed hydrogel network allows the access of light into the scaffold, thereby enabling homogeneous bacteria growth throughout the entire structure.

Bioink transparency is required for efficient light transmission to drive photosynthesis^28^. Light attenuation due to absorption and scattering within the bioink is one of the main challenges to overcome during cultivation of encapsulated photosynthetic species^35^. The F127-based hydrogel transmitted light (76 ± 3%) over the entire visible wavelength range (400–750 nm); the BG11-ASNIII medium used to culture PCC 7002 had 95 ± 4% transmittance in the same range **(Figure 1d)**. The addition of the photoinitiator lithium phenyl-2,4,6-trimethylbenzoylphosphinate (LAP), which absorbs light below 420 nm, reduced transmittance at low wavelengths that are not relevant for PCC 7002 photosynthesis **(Figure S1)**^36^. The encapsulation of cyanobacteria decreased the transmittance of the final construct to 28 ± 8% and 31 ± 9% before and after cross-linking, respectively, due to scattering and productive absorption of light by the photosynthetic bacteria **(Figure S2)**. Overall, the optical properties of the photosynthetic living material were suitable to enable cyanobacteria viability and growth when encapsulated within F127-based hydrogel.

To structure the photosynthetic living materials for enhanced carbon sequestration and longevity, we designed the F127-based hydrogel for both direct ink writing and light-based additive manufacturing. The ink exhibited shear-thinning (shear-thinning index n = 0.09, n < 1) and elastic recovery (∼90%) after high shear, demonstrating its feasibly for extrusion-based printing **(Figure S3 & S4)**. To stabilize printed structures for long-term use, the hydrogel mixture was photo-cross-linked (λ = 405 nm; I = 8 mW cm^-2^; t = 60 s) to increase the final storage modulus (G′) of the construct **(Figure 1c, 1e & S5)**.

### Dual CO_2_ sequestration capacity of photosynthetic living materials

Dual CO_2_ sequestration in the photosynthetic living materials was provided by PCC 7002 biomass growth and insoluble carbonate precipitate formation through MICP. To further investigate both aspects of carbon sequestration, uniform circular samples (V_sample_ = 40 μL, d = 10 mm) were fabricated via direct ink writing and incubated for 30 days **(Figure 2a)**. During the incubation, BG11-ASNIII medium was changed every 5 days. From day 5, the concentration of Ca^2+^ in the medium was set to 8.65 mM via CaCl_2_ addition to simulate natural seawater conditions^37^ and to promote MICP. To highlight the CO_2_ sequestration abilities of the living material, no additional HCO_3_^-^ was dissolved in the medium. As such, all HCO_3_^-^ essential for cell metabolism came from the dissolution of gaseous atmospheric CO_2_.

**Figure 2.**
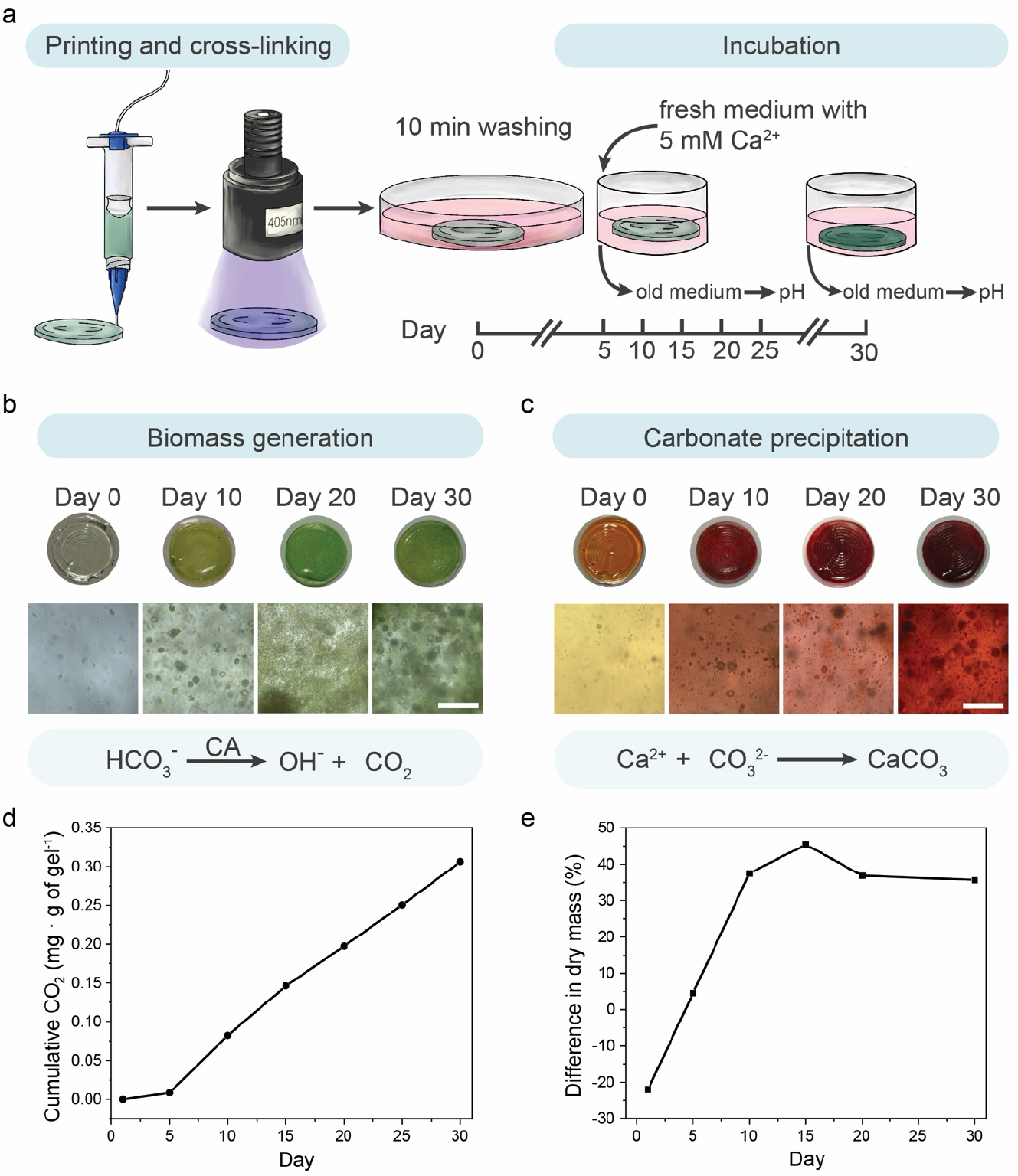
Dual carbon sequestration in photosynthetic living materials. **a)** Standardized discs of the photosynthetic living material were printed and cross-linked prior to incubation in PCC 7002 culture medium for 30 days. Samples were exposed to light (λ = 405 nm, I = 8 mW cm^-2^) and the medium was exchanged every 5 days throughout the incubation. **b)** Biotic samples containing PCC 7002 generated biomass during the 30-day incubation, evidenced by the increasingly green color of the discs and expansion of cell clusters. Biomass accumulation was enabled by carbon utilization via carbonic anhydrase (CA) catalysis of HCO_3_^-^. Scale bar, 100 μm. **c)** In addition to biomass accumulation, carbon sequestration in the form of CaCO_3_ precipitates was visualized by alizarin red staining of the biotic samples over 30 days. Scale bar, 100 μm. **d)** Biotic samples continually sequestered CO_2_ over the 30-day incubation in the form of biomass accumulation (∼0.31 mg CO_2_ per g of gel). **e)** Both biomass growth and CaCO_3_ precipitation contributed to the increase in dry mass of the biotic samples. The cumulative CO_2_ sequestration by biotic samples was reflected by the difference in normalized dry mass between biotic and abiotic samples. Over the incubation period of 30 days, biotic samples accumulated 36% more dry mass than the abiotic samples.

PCC 7002 growth within the living material was observed directly by microscopy and indirectly by measured pH changes in the surrounding medium **(Figure 2b)**. After 10 days of incubation samples exhibited visible clusters of PCC 7002 cells as well as many individual cells throughout the volume of the hydrogel matrix. The number of clusters and individual cells increased with time. The change in pH of the culture medium was attributed to carbonic anhydrase activity in the carboxysome of the bacterial cells. Using HCO_3_^-^ as a substrate, carbonic anhydrase generates equimolar amounts of OH^-^ ions, which are excreted into the surrounding medium, and CO_2_ molecules, which are integrated into bacterial metabolism^17,27^. As the medium was changed every 5 days, the cumulative amount of OH^-^ ions was reflected in the pH increase of the medium from the initial pH value of 6.5. The cumulative OH^-^ ion production served as a surrogate measure of carbonic anhydrase activity and CO_2_ integration into biomass ^38^. Between days 5 and 10, the pH of the biotic samples increased to 9.5, corresponding to approximately three orders of magnitude increase in OH^-^ concentration. In each subsequent 5-day interval, the pH increased by 1.8–2.9 with an average of more than two orders of magnitude increase in OH^-^ concentration **(Figure S6a)**. The cumulative CO_2_ sequestered was calculated based on the pH change of the medium **(calculations in the Supplementary Information, CO**_**2**_ **sequestration by biomass growth (pH change) in living materials)**. Over the first 15 days the cumulative amount of CO_2_ sequestered per gram of living material was 3.3 μmol (0.15 mg) and reached a value of 7 μmol (0.31 mg) by the end of the 30-day experiment, an increase that reflects CO_2_ sequestration corresponding to biomass accumulation **(Figure 2d)**. The pH did not change in abiotic control samples throughout the incubation **(Figure S6b)**.

To confirm CO_2_ sequestration via the production of insoluble carbonate precipitates via MICP, precipitates were visualized using calcium staining. In the presence of divalent ions such as Ca^2+^ and Mg^2+^, cyanobacteria go through the biomineralization process to form gypsum, calcite, and magnesite ^39^. We confirmed the accumulation of Ca^2+^ in the hydrogels using Alizarin red staining **(Figure 2c)**. On day 0, abiotic and biotic samples were mainly orange after staining with green PCC 7002 cells visible in the biotic samples, indicating minimal Ca^2+^ accumulation initially. The abiotic samples remained orange throughout the 30-day incubation **(Figure S7)**. Biotic samples stained red by day 10, indicating Ca^2+^ accumulation in the whole volume of the hydrogel and the dark red color was observed around growing bacteria clusters over time. To assess the amount of biomass and precipitate formation, the mass of abiotic and biotic samples was measured and compared. Over the incubation period of 30 days, the biotic samples had approximately 36% more dry mass than control abiotic samples **(Figure 2e)**. In total, biomass and carbonate precipitates in biotic samples accounted for approximately 45% of the final sample mass **(calculations in the Supplementary Information, total equivalent CO**_**2**_ **sequestration calculation in living materials)**.

To quantify the extent of precipitate generation, the organic biomass and polymer matrix were removed by thermal decomposition. As the temperature (T = 600 °C) was above the decomposition temperature of organic matter (polymer and biomass) and below the decomposition temperature of carbonate compounds ^40^, only the insoluble carbonate precipitates remained after thermal decomposition. The mass of the inorganic precipitates corresponded to 50 μmol (2.2 ± 0.9 mg) of CO_2_ sequestered via MICP per gram of hydrogel **(Figure S8 and calculations in the Supplementary Information, total equivalent CO2 sequestration calculation in living materials)**. CO_2_ sequestration via MICP was approximately one order of magnitude more than that achieved via biomass growth, making the dual CO_2_ sequestration a more efficient carbon sink, and indicating that the majority of the sequestered CO_2_ had been stored in a more stable mineral form.

### Composite materials form during the life cycle

Having demonstrated that photosynthetic living materials are capable of CO_2_ sequestration via both biomass accumulation and carbonate formation, we further investigated the composition of the mineral phase produced during incubation. MICP results in the formation of calcium carbonates that vary in stability and solubility with calcite being the most stable polymorph and, therefore, desirable for carbon sinking purposes **(Figure 3a)**^41^. XRD analysis of precipitates in biotic samples revealed a highly crystalline structure with the main peaks corresponding to calcite structure with a noticeable shift of the main diffraction peak representing the (104) plane **(Figure 3b)**. The observed shift likely indicated compositional variation due to the incorporation of magnesium in the carbonate structure as previously observed in marine environments and, in this case, due to the use of simulated seawater (ASNIII medium)^42^. The XRD was obtained after thermal decomposition (T = 600 °C, which is below the decomposition temperature of CaCO3)^40^ as the signal from the F127-based matrix was much stronger than the one of the precipitates **(Figure S9)**.

**Figure 3.**
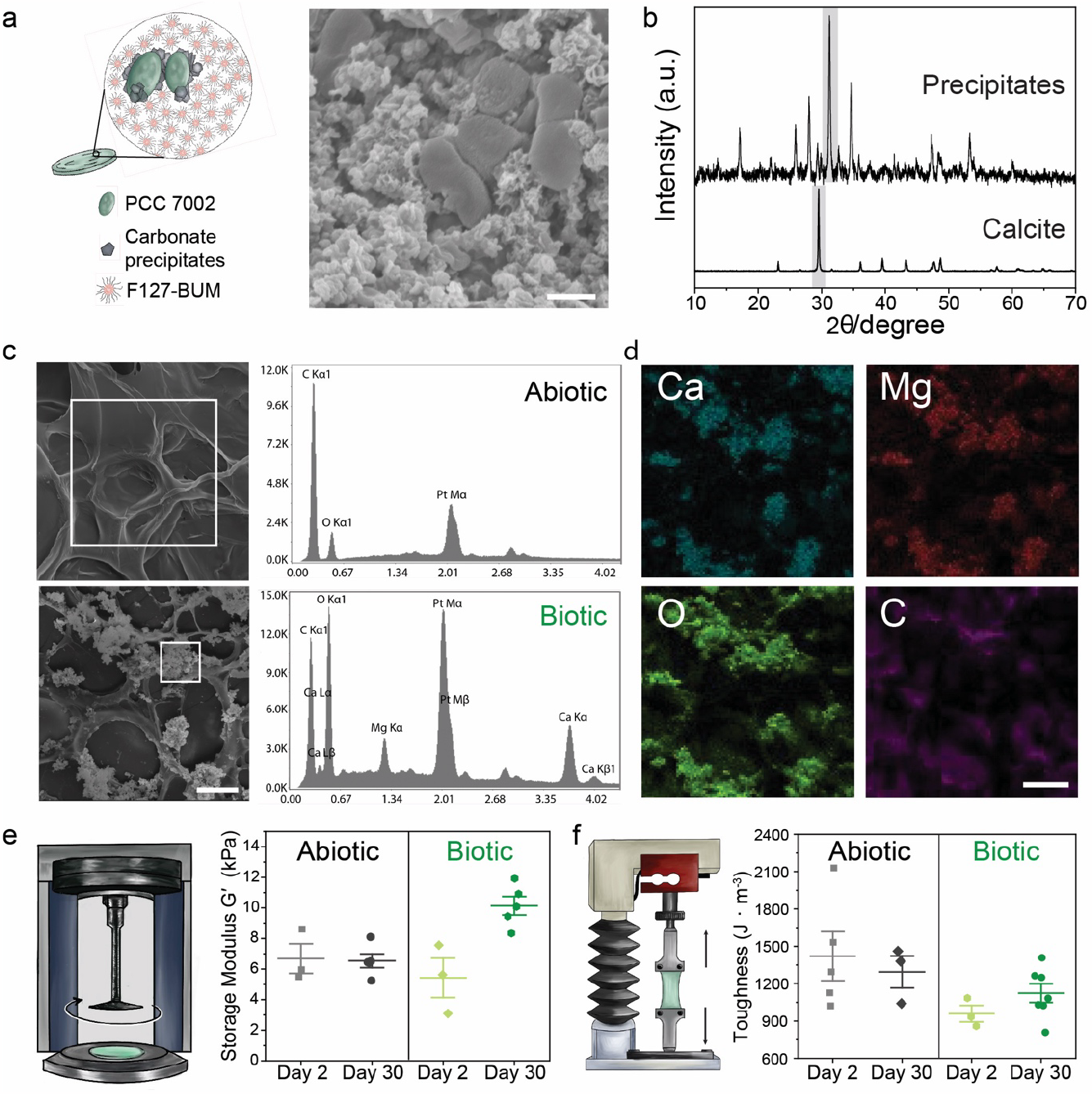
Carbonate formation within photosynthetic living materials. **a)** During incubation, carbonate precipitates formed around PCC 7002 cells in the photosynthetic living materials as visualized by SEM. Scale bar, 1 μm. **b)** XRD analysis of the remaining precipitates, following removal of the polymer content and bacterial biomass by thermal decomposition, indicated the presence of carbonates in the biotic samples ^48^. **c)** EDS analysis on the selected region indicated in the SEM images of abiotic and biotic samples corroborated the presence of calcium and magnesium containing carbonates only in the biotic samples. Scale bar, 20 μm. **d)** EDS elemental mapping indicated co-localization of Ca, Mg, O, and C atoms in the mineralized regions of the biotic samples. Scale bar, 20 μm. **e)** The storage modulus (G ′), measured by shear rheometry, increased during the 30-day incubation in the biotic samples. The modulus of abiotic samples remained constant. **f)** Correspondingly, the toughness of the abiotic samples did not change substantially between day 2 and day 30, while the toughness of biotic samples increased during the 30-day incubation.

Precipitates distributed throughout the polymeric matrix **(Figure 3c)**. Energy dispersive X-Ray analysis (EDS) indicated mainly carbon and oxygen in the abiotic matrix whereas peaks of calcium and magnesium were prominent in the areas with precipitates in biotic samples. Elemental mapping corroborated these data with calcium, magnesium, carbon, and oxygen in overlapping regions near clusters of cyanobacteria, suggesting the pericellular formation of carbonates **(Figure 3d)**.

The formation of precipitates in hydrogel matrices via MICP can reinforce the mechanical properties of the material^17,43,44^. The storage moduli (G′) of the printed biotic samples on day 2 of the experiment were slightly lower compared with abiotic samples (5.4 ± 2 kPa and 6.7 ± 2 kPa, respectively). Over the course of 30 days, the modulus increased to 10.1 ± 1 kPa in the biotic case while it did not change for abiotic samples (6.5 ± 1 kPa) **(Figure 3e)**. Comparable results were obtained for the Young’s moduli (E) of the samples using tensile tests of cast samples **(Figure S10)**. We associated the increase in modulus with the formation of reinforcing inorganic precipitates^45^, though increases in biomass could also contribute to the change in Young’s moduli^46,47^. A similar trend was observed for the material toughness. After 30 days, the toughness of the biotic samples increased compared to day 2 (from 960 ± 110 J m^-3^ to 1120 ± 200 J m^-3^) **(Figure 3f)**.

### 3D-printed photosynthetic living structures for dual carbon sequestration

While simple discs served as a standardized shape for the systemic characterization of photosynthetic living materials, the geometry of the living structures was tailored via digital design and fabrication to improve the CO_2_ sequestration and long-term viability of the photosynthetic living materials. We designed 3D lattice structures with strut sizes between 0.15 mm and 0.70 mm to facilitate gas and nutrient transport within the printed constructs. The transport of gases and liquid medium as well as access to light are essential for the efficient functioning of biologic processes (including carbon sinking) in the photosynthetic living materials^9,31^. Drawing inspiration from cellular fluidics^32^, we employed a lattice design to engineer structures whereby growth medium was passively transported vertically through the construct via capillary forces **(Figure 4a)**. In this manner, the structure did not need to be fully immersed in the simulated seawater, minimizing medium use, and facilitating gas transport from the surrounding air to the living material. To achieve high resolution and defined internal porosity, light-based volumetric 3D printing was used to fabricate the lattice structures. Centimeter-scale objects were printed with complex geometries and an optical resolution of 28 x 28 μm within tens of seconds **(Figure 4b)**. The 3D-printed photosynthetic living structure remained viable for over a year, during which it was continuously performing dual carbon sequestration. After 30 days of incubation, the structure was able to stand vertically on a flat surface and liquid was actively drawn up via the construct’s internal structure. The printed sample stiffened further over prolonged incubation due to the accumulation of carbonate precipitates. This mechanical enhancement was observable through the structure’s increased capacity to maintain an upright position at day 30, 60,120 and 365 and a final modulus of 111 ± 7 kPa (**Table S3**). Over the incubation period of 400 days the living structures sequestered 26 ± 7 mg of CO_2_ per gram of hydrogel material in the form of carbonate precipitates **(Table S4)**. This prolonged viability of the photosynthetic living materials showed that dual carbon sequestration can take place beyond 30 days, especially within rationally designed 3D structures.

**Figure 4.**
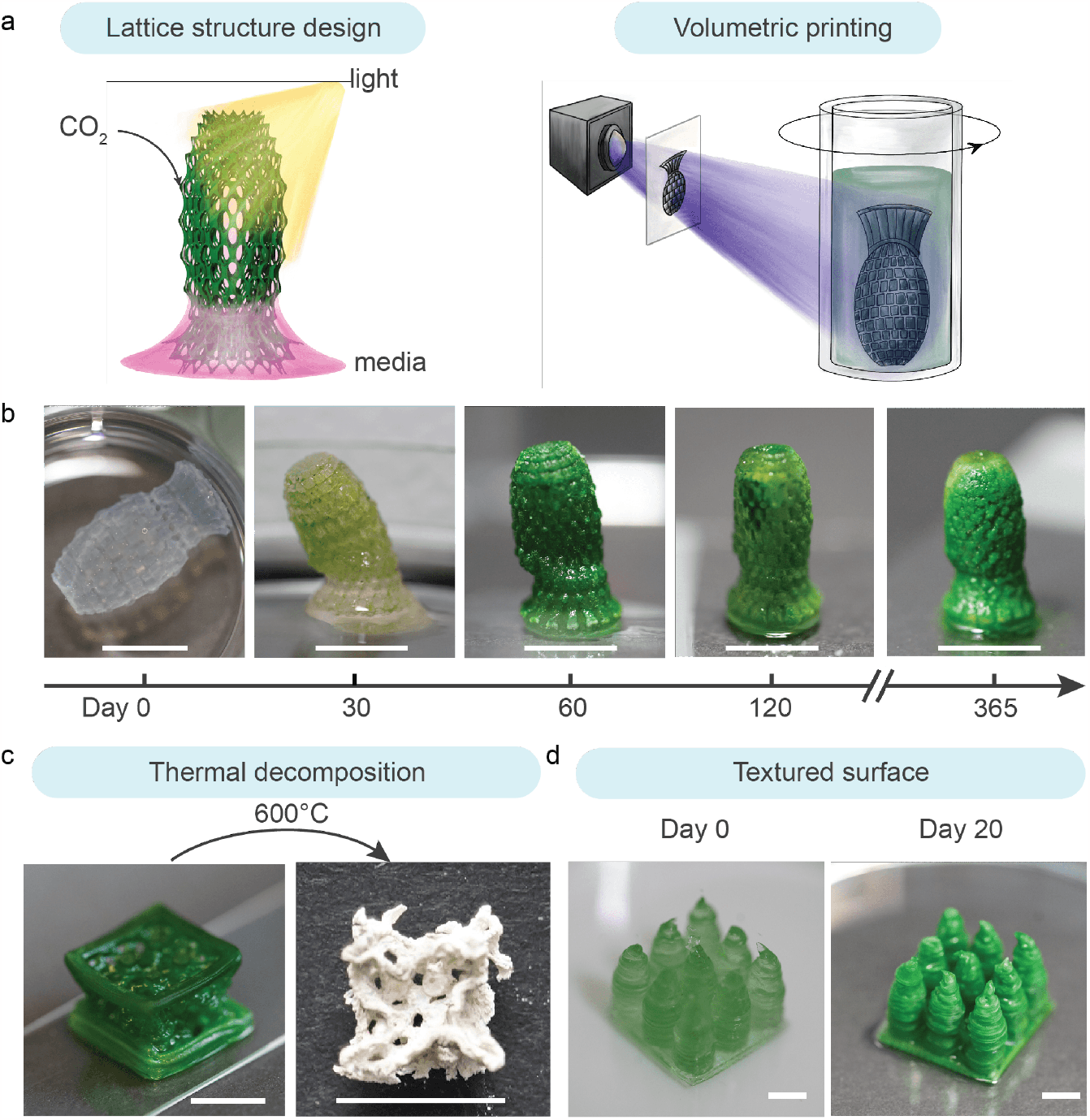
Digital fabrication of photosynthetic living structures for dual carbon sequestration. **a)** Lattice structures were digitally designed to facilitate light exposure and medium transport via passive capillary wetting, enabling growth of the photosynthetic living materials. The structures were fabricated using volumetric printing **b)** The 3D-printed lattice evolved during the incubation from a free-floating structure at day 0 to a self-standing and partially mineralized construct at day 30. The photosynthetic living material continued to evolve over time and exhibited robust viability up to 365 days. Scale bar, 1 cm. **c)** A larger cubic lattice structure was printed using direct ink writing and incubated for 60 days. The sample was thermally decomposed (T = 600 °C) to remove the biomass and polymer matrix, revealing a templated mineral construct. Scale bar, 1 cm. **d)** To increase the volume of viable material per unit of surface area, a coral-inspired structure was printed via direct ink writing (day 0), which demonstrated improved growth (day 20) as compared with a flat construct of the same surface coverage (footprint) and volume owing to enhanced light exposure. Scale bar, 0.5 cm.

To demonstrate the scalable production of photosynthetic living materials, direct ink writing was used to print similar lattice structures at a larger scale. A body-centered cubic (BCC) lattice structure (unit cell length l = 5 mm, strut diameter d = 0.12 mm) was printed. The structure passively perfused medium to the top surface while only partially immersed in BG11-ASNIII medium (**Figure S12**). After 60 days of incubation, the lattice was thermally decomposed (T = 600 °C). The remaining carbonate precipitates after thermal decomposition retained the shape of the porous lattice structure **(Figure 4c)**, indicating that carbon sequestration was homogeneous throughout the structure despite being partially immersed in medium.

This living material also has the potential to be applied on existing surfaces as coatings for dual carbon sequestration. Starting from bulk material, we observed 5 mm to be an optimal material thickness to maintain PCC 7002 viability (0.5 g_material_ cm^-2^) **(Figure S13)**^9^. To maximize the viable volume of the material per surface area, textured surface was designed as 3 x 3 pillar array on a 2 x 2 cm base to minimize self-shielding of light similarly to coral reef structures^49^. With this design we increased the printed gel volume by 150% while maintaining the same 1 cm^2^ footprint (to 0.75 g_material_ cm^-2^) and without compromising bacteria viability compared to a bulk hydrogel **(Figure 4d, S14)**. This result further highlights how the synergy between living materials and the design of living structures can increase the efficiency of dual carbon sequestration.

### Discussion

The design of our photosynthetic living materials builds upon many recent advances in engineered living materials^9,13,50,51^. The diverse functionality of microorganisms has enabled the design and application of engineered living materials in various fields. For instance, bacteria-based biosensors are used to detect target molecules, such as glucose or specific pathogens^52,53^. Living skin patches adopt skin microbiomes to control bacterial outgrowth on the wounded area^54^. Utilizing the bacterial-mineralization process, microorganism-based living building materials can self-heal and regenerate^16,17,27^. Living materials have also been used to replace conventionally pollution-intensive processes, such as applications in the energy industry, the production of textiles or structural materials, due to their inherent bioremediation abilities, especially through the carbon sequestration capabilities of certain microorganisms^55–59^.

Building on these concepts, we engineered printable photosynthetic living materials that were capable of carbon sequestration through biomass accumulation and inorganic carbonate precipitation. To enhance the carbon sequestration potential of our photosynthetic living material, we used different additive manufacturing approaches to design biomimetic porous structures that drew upon principles outlined by cellular fluidics^32^. Tailoring construct geometry via additive manufacturing enabled our photosynthetic living structures to survive beyond one year while continuously undergoing dual carbon sinking. Throughout the incubation period of 400 days, our photosynthetic living material continuously performed dual carbon sequestration, with most of the sequestered carbon stored in the stable mineral form. The total amount of CO_2_ sequestered through mineral formation after 400 days was 26 ± 7 mg per gram of photosynthetic living material. This amounts to approximately 12 times more than the CO_2_ sequestered through mineral formation during our 30-day experiments (2.2 ± 0.9 mg per gram of living material). With minimum requirements of sunlight and atmospheric CO_2_, our photosynthetic living materials showed a consistent carbon sequestration efficiency throughout the entire incubation period, and demonstrated the potential of using natural carbon sequestration processes at scale^60^.

As a comparison of the efficiency of this living approach, similar CO_2_ capture initiatives via chemical mineralization, such as carbonating recycled concrete aggregates, have been able to sequester CO_2_ at a quantity of 6.7 mg per gram of recycled concrete aggregates^61^. Thus, our first-generation photosynthetic living material (26 ± 7 mg of CO_2_ sequestration per gram of living material) may be competitive with industrial chemical mineralization process. Moreover, the carbon sequestration capacity of our photosynthetic living materials is not limited by the availability of essential chemicals, such as Ca(OH)_2_ availability in recycled concrete aggregates. Comparatively, CO_2_ sequestration via chemical and biological mineralization is less efficient than carbon capture and storage (CCS); however, CCS requires a concentrated CO_2_ source as well as controlled temperature and pressure to operate. Our photosynthetic living materials function at ambient conditions with atmospheric CO_2_ as the sole carbon source, which highlights the ability to use photosynthetic living materials as a complementary approach in a more distributed manner.

Implementing photosynthetic living materials in broad application still requires improved usability and upscaling of the material production. This can be achieved by taken advantage of recent advances in biofabrication for the creation of larger scale porous^62^ or granular^63^ structures. In addition, methods to engineer optical structures at scale for efficient light harnessing, may further improve efficiency^49^. Photosynthetic living materials also hold promise for future applications as surface coatings for green building materials or bioreactors in commercial-scale sequestration plants, enabling bioremediation of CO_2_ emissions and supporting the realization of carbon-neutral to carbon negative infrastructure. Their simple requirements and easy maintenance also enable possible installation in various environments, ranging from urban to rural landscapes, for long-term and sustainable CO_2_ sequestration. To further enhance the efficiency of the system, we foresee the possibilities to genetically modify or select microorganisms or microorganism consortia with higher photosynthetic rates.

## Conclusion

We have introduced a new cyanobacteria-laden photosynthetic living material for dual carbon sequestration, achieved through the combination of simultaneous biomass generation and the formation of insoluble carbonate precipitates. The photosynthetic living materials performed dual carbon sequestration over an extended life of up to one year with light and atmospheric carbon as its energy and carbon source. The base formulation of this living material enabled processing via direct ink writing as well as light based additive manufacturing. As a result, the final design of living structures was tailored for enhanced carbon sinking via increased surface area and porosity. Furthermore, the spatiotemporal control provided by 3D printing allows scale up for potential applications in disparate fields, such as civil engineering and architecture.

## Acknowledgements

This work was done within the framework of the ALIVE initiative (Advanced Engineering with Living Materials) and funded by the SFA-AM program (Strategic Focus Area – Advanced Manufacturing) by the ETH Board. This work was also supported by the Swiss National Science Foundation (200021_184697; MWT) and the NIH-funded Wyoming IDeA Networks of Biomedical Research Excellence Program (P20GM103432; JSO).

## Experimental Section

### Bacteria culture and encapsulation

The cyanobacteria *Synechococcus* sp. PCC 7002 (Pasteur Institute, France) were cultured in BG11-ASNIII medium mixture **(described in ESI, cyanobacteria culture conditions, and Table S1)** in Erlenmeyer flasks on a shaker plate with constant shaking (150 rpm) under full spectrum white light (180 μmol photons m^-2^ s^-1^) with a 12-hour day (on)/night (off) cycle. Optical density of the bacteria suspension at 730 nm (OD_730nm_) was measured using UV-visible light spectrophotometer (Lambda 35, Perkin Elmer) to monitor bacterial culture growth phases. For encapsulation, cyanobacteria were collected at the early exponential growth phase.

### Bioink preparation

Stock solutions of 18 wt% F127 in culturing medium, 30 wt% of F127-BUM **(synthesis described in ESI, F127-bis-urethane methacrylate synthesis)** in Milli-Q water, and 5 wt% LAP **(synthesis described in ESI, LAP synthesis)** in Milli-Q water were mixed at 3:1:0.082 ratio to reach final concentrations of 13.2 wt% F127, 7.3 wt% F127-BUM, and 0.1 wt% LAP. For the bioink preparation, cyanobacteria suspension was centrifuged at 3500 rcf for 5 mins, the supernatant was discarded, and the cell pellet was resuspended in the prepared bioink mixture to reach an equivalent optical density OD_730nm_ = 0.8. For volumetric printing, an equivalent optical density OD_730nm_ = 0.3 was used. The equivalent OD_730nm_ calculation is described in **Equation S1**.

### Bioink characterization

The bioink was characterized using strain-controlled shear rheometer (MCR 502; Anton-Paar). The samples were loaded on a glass bottom plate with a light source (λ = 405 nm, I = 8 mW cm^−2^; M405L3, Thorlabs) underneath and measured using 20 mm plate-plate probe with 0.8 mm gap size at 25 °C. Cross-linking kinetics of the abiotic gels was evaluated by using oscillatory time sweep (ω = 10 rad s^-1^; γ = 0.1%). The samples were irradiated after 60 s of initial oscillation to induce photopolymerization of F127-BUM in the gel mixture until the plateau storage modulus G′ was reached.

UV-vis spectra of the medium and hydrogel samples were measured using a plastic cuvette (1 cm light path) with a UV-visible light spectrophotometer (Lambda 35, Perkin Elmer). The transmittance was measured over the range of 800-400 nm.

### Biofabrication

3D samples for the measurements were fabricated using a pneumatic-driven direct ink writing 3D printer (BioX, Cellink). The bioink was loaded into 3 mL cartridges with a 22G conical nozzle (Ø = 0.41 mm) and the cartridge was heated to 37 °C for 5 mins prior to printing. The printing was performed at a pressure range of P = 40-60 Pa and a speed of v = 5-10 mm s^-1^. Single layer circular disc-shaped samples (Ø = 1 cm, h = 150-350 μm) were modelled in Rhinoceros 3D and G-code generation was done via Slic3r. The more complex lattice structures were designed with Rhinocerous 3D (Rhino7) and Grasshopper software. The printed disc-shaped samples were cross-linked for 1 min using an LED light source (λ = 405 nm light, I = 8 mW cm-2, Thorlabs GmbH, Germany), followed by a 10 min washing step in BG11-ASNIII medium. The washing medium was discarded, and fresh medium was added to the samples for bacteria growth.

For volumetric printing, a Tomolite tomographic bioprinter (Readily3D, Switzerland) was used as described elsewhere ^64,65^. All stl. models were generated with Rhino7 and Grasshopper software. The size constraints for the Tomolite vials were up to 25 mm in height and 14 mm in diameter. The lattice shown in **Figures 4a and 4b**, is 21.85 mm tall and 11.85 mm in diameter, with a volume of 101.87 mm^3^ and a surface area of 2145.84 mm^2^. The lattice is radially arrayed into 16 sectors, in 3 concentric rings, and 20 vertical segments. This generated a lattice with 960 pores, with pore diameter from 0.5-1.0 mm, and strut diameters from 0.15-0.70 mm (median diameter 0.20-0.24 mm). The lattices were sliced into tomographical projection planes using a commercial software (Apparite, Readily3D, Switzerland). Bioink was loaded into glass printing vials (20 mm outer diameter) and solidified at 25°C for 5 mins before printing. Cross-linking of the bioink was induced by a light source with a 405 nm wavelength with a pre-calibrated light dosage of 94 mJ cm^-2^. The averaged light intensity during the printing process was set to be 8 mW cm^-2^. After printing, the vial was cooled to 4 *°*C to liquify the unpolymerized bioink. The retrieved polymerized structure was immersed in culturing medium to allow removal of excess LAP and residual bioink. The sample was incubated beyond one year with the above-mentioned culturing conditions.

### Living material incubation conditions

The printed circular disk-shaped samples were cross-linked for 1 min using an LED light source (λ = 405 nm light, I = 8 mW cm^-2^, Thorlabs GmbH, Germany), followed by a 10 min washing step in BG11-ASNIII medium. After that, printed samples were incubated under the same light conditions as liquid culture. The samples were incubated in BG11-ASNIII medium for 5 days with a change to fresh medium on day 2. The old medium was collected at each change for pH measurements. Fresh BG11-ASNIII medium with an additional 5 mM CaCl_2_ was added as the calcium-rich culturing medium. The medium was changed and collected every 5 days (on day 5, 10, 15, 20, and 25) until day 30. The collected medium was used for pH measurements to evaluate total sequestered CO_2_ as fully **described in the Supplementary Information (CO**_**2**_ **sequestration by biomass growth (pH change) in living materials)**. The samples were inspected using optical microscope (Panthera Classic, Motic) equipped with Plan UC 10x/0.25, 40x/0.65 Ph2 and 100x/1.25 oil objectives and camera (Moticam S, Motic). The images were captured with Motic Images Plus 3.090 software.

### Biomass characterization

5 of the freshly printed (day 0) abiotic and biotic disc samples were washed in Milli-Q water for 10 min to remove salt deposited on the surface from the culturing medium. The samples were then dried under a constant laminar flow in a 1.5 mL microcentrifuge tube of known mass to evaporate the water entrapped within the material. The dried samples together with the microcentrifuge tube were weighed and the mass was used as the initial benchmark for characterization.

On day 5, 10, 20 and 30, five of the abiotic and biotic samples were randomly selected and washed with the same procedure. The samples were subsequently dried under the same condition before weighing. The mass was normalized to the average mass of the abiotic or biotic discs on day 0.

### Inorganic precipitate characterization

Carbonate precipitate deposits in circular disc-shaped hydrogels were stained using 40 mM Alizarin red S solution in Milli-Q water. Each sample was fully immersed in 2 mL of Alizarin red S solution for 20 s. The sample was then washed in 200 mL of Milli-Q water 3 times for 5 min and imaged using an optical microscope.

The extent of carbonate precipitate formation within abiotic and biotic samples was characterized by thermal decomposition at atmospheric pressure. Five dry abiotic or biotic samples were loaded into a porcelain crucible and heated to 600 °C (ramp rate = 2 °C min^-1^) for 2 hours. The samples were then cooled to room temperature and the remaining mass was measured. In addition, the remaining precipitates were collected for X-ray diffraction (XRD) analysis with a PANalytical Empyrean diffractometer (Cu Kα radiation,45 kV and 40 mA) with an X’Celerator Scientic ultrafast line detector and Bragg-Brentano HD incident beam optics. Samples were measured over the 2θ range 10-90° for 1 h with a step increment of 0.016°. Results were compared with reference patterns from the ICDD database.

Lyophilized samples **(procedure described in the Supplementary Information, sample lyophilization)** were used for Scanning Electron Microscope (SEM) characterization. Samples were sputtered with 10 nm of Pt/Pd 80/20 with a metal sputter coater (CCU-010, Safematic, Zizers, Switzerland). SEM imaging was done using a secondary electron detector with a scanning electron microscope JEOL JSM-7100F (accelerating voltage V = 12 kV, working distance WD = 10 mm) equipped with an Ametek-EDAX EDS detector (Si(Li) 20 mm). Element analysis was performed with the EDAX Team software.

### Living material mechanical properties

Viscoelastic properties of the printed hydrogels at different time points during incubation were evaluated using the same rheometer as for the bioink characterization equipped with a Peltier stage and 8 mm sandblasted probe (gap size h = 0.15-0.35 mm, normal force F_N_ = 0.1 N) at 25 °C. Samples were cut from disc-shaped samples using a metal punch (Ø = 8 mm) to match the rheometer probe geometry. Dynamic oscillatory frequency sweep measurements were performed in the range of 100 to 0.1 rad s^-1^ (γ = 0.3%). Young’s modulus was calculated as E = 2G (1 + ?), using storage value G′ at ω = 10 rad s^-1^ and an assumption for Poisson ratio of ? = 0.5.

To obtain homogeneous samples for tensile testing, abiotic and biotic bioink were molded into a rectangular-shaped mold (12 mm × 50 mm × 1 mm, width x length x thickness) and cross-linked for 2 mins (λ = 405 nm light, I = 8 mW cm^-2^). Samples were then washed and incubated under the same conditions as the disc-shaped samples. Uniaxial tensile testing was performed on the samples incubated for 2 and 30 days using a tensile testing machine (Stable Micro Systems) equipped with two parallel metal screw-clamps. A 2 mm notch was cut with a razor blade on one side of the sample prior to the stress-strain measurement. The sample was placed securely between the two parallel clamps with an initial grip-to-grip distance of 300 mm. Stress-strain measurement was performed with a constant stretch speed of 0.5 mm s^-1^ until failure. The sample toughness was calculated by measuring the area under the stress–strain curve and Young’s modulus of the sample was calculated as the slope of the linear regime of the curve **(Figure S11**).

Compressive tests were conducted on the same tensile testing machine (Stable Micro Systems) equipped with a steel probe (Ø = 0.2 mm). The compression test was done at a constant deformation rate of 0.05 mm min^−1^ on a cylinder-shaped photosynthetic living material sample after 400 days of incubation (Ø = 10 mm, h = 3 mm). The compressive modulus was obtained by measuring the slope of the linear regime of the stress–strain curve **(Table S3)**.

