## Supplementary Information for "Dual carbon sequestration with photosynthetic living materials"

Figure S9. XRD analysis of the abiotic and biotic samples after 30 days of incubation. The samples were first lyophilized to remove any water content, followed by homogenizing in 1 mL of absolute ethanol for 2 minutes. The major peaks at 18 and 23 correspond to the Pluronic F127 micelles and the calcite peaks are not prominent due to the strong signal from the F127 micelles. (a) XRD spectra over 2θ of 10 - 70°, (b) zoomed-in XRD spectra between a 2θ of 25 and 70°. The signal from the F127-based matrix was much stronger than the one of the precipitates and the signal at 31° could be associated with the calcite peak post thermal decomposition. ........ 17

**List of materials**

Pluronic F127 (P2443); anhydrous dichloromethane (DCM anhydrous, 900633); dibutyltin dilaurate ($C_{32}H_{64}O_4Sn$, 291234); 2-isocyanatoethyl methacrylate ($C_7H_9NO_3$, 477060); sodium chloride (NaCl, 71380); magnesium sulfate anhydrous ($MgSO_4$, M7506); sodium nitrate ($NaNO_3$, 71755); magnesium chloride hexahydrate ($MgCl_2 \cdot 6H_2O$, 63068); citric acid ($C_6H_8O_7$, C0759); sodium carbonate ($Na_2CO_3$, 13418); ferric ammonium citrate ($C_6H_{5+4y}Fe_xN_yO_7$ , F5897); BG11 broth (73816); A5+ Co Trace metals (92949); vitamin $B_{12}$ (V2876); EDTA disodium magnesium ($(NaOOCCH_2)_2NCH_2CH_2N(CH_2COO)_2Mg \cdot xH_2O$, 317810); alizarin red S ($C_{14}H_7NaO_7S$, A5533); lithium bromide (LiBr, 213225), dimethyl phenylphosphonite ($C_6H_5P(OCH_3)_2$, 149470); diethyl ether (32203); 2,4,6-trimethylbenzoyl chloride (682519-5G); 2-butanone ($C_2H_5COCH_3$, 360473) were purchased from Sigma-Aldrich (Germany). Potassium chloride (KCl, 26764.232); methanol ($CH_3OH$, 02000347); dipotassium phosphate anhydrous ($K_2HPO_4$ anhydrous, 26931.263) were purchased from Thermo Fisher Scientific (Switzerland). Anhydrous calcium chloride ($CaCl_2$ anhydrous, 349615000) was purchased from Acros Organics.

**F127-bis-urethane methacrylate synthesis**

F127-bis-urethane methacrylate (F127-BUM) was synthesized by adapting the previously reported method by Millik et al. [1]. Pluronic F127 (m = 30 g) was vacuum dried for 2 h and then dissolved in anhydrous DCM (V = 275 mL) by magnetic stirring at 30 °C until Pluronic F127 was fully dissolved. 6 drops of dibutyltin dilaurate was then added. A mixture of 2-isocyanatoethyl methacrylate (V = 1.75 mL) and DCM (V = 25 mL) was added to the Pluronic F127 solution at a rate of 1 drop per second and the addition process was allowed to proceed under argon atmosphere at 30 °C overnight.

The reaction was then quenched with methanol (V = 30 mL), concentrated by a rotatory evaporator at 45 °C, and subsequently precipitated in 800 mL of ice cold diethyl ether. The reaction product was recovered by centrifugation (at 3000 rcf for 10 min at 4 °C) followed by two additional washing with diethyl ether and centrifugation steps. Synthesized F127-BUM was then dried under vacuum and stored at -20 °C until use. Pluronic F127 modification was confirmed via $^1H$ NMR (400 MHz, CDCl3): δ 6.06 (m, 2 H), 5.54 (m, 2 H), 5.15 (m, 2 H), 4.16 (m, 8 H), 3.59 (m, 770 H), 3.49 (m, 117 H), 3.35 (m, 56 H), 1.89 (s, 6z H), 1.08 (m, 170 H), (**Figure S15**).

**LAP synthesis**

Photoinitiator lithium phenyl-2,4,6-trimethylbenzoylphosphinate (LAP) synthesis was adapted from previously reported method [2,3]. 2,4,6-trimethylbenzoyl chloride (V = 2.9 mL) was slowly added to dimethyl phenylphosphonite (V = 2.8 mL) under argon. The mixture was stirred overnight at 25°C. Lithium bromide (m = 6.1 g) was dissolved in 2-butanone (V = 100 mL) and added to the reaction flask. The temperature of the mixture was then increased to 50 °C for 10 mins to facilitate precipitate formation. The precipitates were recovered using filtration and washed 3 times with 2-butanone. LAP was then dried under vacuum and stored at -20 °C until use. Product formation was confirmed via $^1H$ NMR (400 MHz, D2O): δ 7.75 (m, 2 H), 7.60 (m, 1 H), 7.51 (m, 2 H), 6.93 (s, 2 H), 2.28 (s, 3 H), 2.06 (s, 6 H) (**Figure S16**).

**Bioink printability characterization**

Bioink printability was characterized using strain-controlled shear rheometer (MCR 502; Anton-Paar). The samples were loaded on a temperature-controlled Peltier plate and measured using a 20 mm plate-plate geometry with 0.8 mm gap size at 25 °C. The self-healing properties of the Pluronic F127 based inks were evaluated using dynamic oscillatory time sweep tests with alternating low strain ($\omega$ = 10 rad $s^{-1}$; $\gamma$ = 0.3%, for t = 120 s) and high strain ($\omega$ = 10 rad $s^{-1}$; $\gamma$ = 1000%, for t = 240 s). The recovery of storage modulus G′ was evaluated by calculating the ratio between initial G′ value and that of G′ after plateau was reached in a low strain interval.

Rotational shear rate measurements were performed in the shear rate range of dγ/dt = 0.1-100 $s^{-1}$. Shear-thinning index, n, and consistency index, K of the bioink were obtained by fitting the viscosity dependance on shear rate to Power-law model: $\eta = K\,(d\gamma/dt)^{n-1}$ (**Figure S4, Table S2**).

**Bacteria culturing conditions**

Culture medium conditions in a mixture of BG11 and ASNIII medium were adapted from the guidelines of the supplier. ASNIII medium was prepared by mixing all components (**Table S1**) in deionized water, then autoclaved and supplemented with A5+ Co Trace metals (1x from initial 1000x stock solution) and vitamin $B_{12}$ (0.010 g $L^{-1}$). ASNIII medium was used at 2x final concentration. BG11 medium was used as received (100x stock solution) at a 2x concentration.

The bacteria culture was cultivated in Erlenmeyer flasks on a shaker plate (150 rpm) at 30 °C and illumination of approximately 180 μmol photons $m^{-2}$ $s^{-1}$ using 12-hour day (on)/night (off) cycle.

**Biotic bioink $OD_{730}$ calculation**

In order to achieve an equivalent optical density of 0.8 at $OD_{730nm}$ in the bioink, the $OD_{730nm}$ of cyanobacteria suspension prior to encapsulation ($OD_{730nm,culture}$) was measured using UV-visible light spectrophotometer (Lambda 35, Perkin Elmer). $OD_{730nm}$ of pure culturing medium ($OD_{730nm,medium}$) was used as a blank. For every 1 mL of bioink required, the volume of initial cell culture required was calculated as described in **Equation S1.**

$$V_{cell\ culture} = \frac{OD_{730\mathrm{nm},culture} - OD_{730\mathrm{nm},media}}{0.8} \times 1\ mL \qquad \text{Equation S1}$$

The growth curve of cyanobacteria culture in BG11-ASNIII medium was obtained by measuring the optical density at 730 nm over a period of 36 days and the data is plotted in **Figure S17.** Absorbance of pure culturing medium was also measured as a background reference.

**Sample lyophilization**

Cyanobacteria-laden (biotic) samples and abiotic samples were first sterilized in 70% ethanol for 1 h. Ethanol was then discarded and 100 mL of Milli-Q water was added to each sample for 10 min to wash any remaining salt deposited on the surface of the samples from the culture medium. After washing, the samples were frozen (T = -20 °C) overnight with fully immersed in Milli-Q water.

A Lyovapor L-300 lyophilizer (BUCHI Labortechnik AG, Flawil, Switzerland) was used with a set pressure of 0.150 mbar and an ice condenser temperature of approximately -101 °C to freeze dry the samples.

**XRD spectra analysis on baseline removal**

The obtained XRD data was imported into Origin (OriginLab Corporation) for baseline removal. To define the baseline, 40 user-defined points with $2^{nd}$ derivative anchoring points method were chosen to match the baseline of the plotted $2\theta$ against intensity curve. The defined baseline was subtracted from the original intensity values and the curve was plotted again with the baseline subtracted.

**Bioink preparation for volumetric printing**

As described bioink was chilled to 4 °C to reach a liquid state before filling into an autoclaved printing vial (outer diameter Ø = 20.0 mm, inner diameter Ø = 15.0 mm). The vial was then heated up to room temperature (T = 25°C) for 5 mins before inserting into the bioprinter.

**Light dosage calibration for volumetric printer**

Light dose testing was performed as per the manufacturer's instructions. 1 mL of bioink at 4 ℃ was loaded into a calibration quartz glass cuvette (Thorlabs, CV10Q14). Light dosage varying from 360 to 40 mW $cm^{-2}$ was applied at different locations of the cuvette to examine the polymerization efficiency. A second cuvette was then loaded, and the same calibration process was done with a narrower range of light dosages. The final optimal light dosage was calculated by an Apparite software (Readily3D, Switzerland).

**$CO_2$ sequestration by biomass growth (pH change) in living materials**

The change in pH of the culture medium was attributed to carbonic anhydrase activity in the carboxysome of the cells which converts $HCO_3^-$ into equimolar amounts of $OH^-$ ions which go into the surrounding medium and $CO_2$ molecules which are integrated into the bacteria metabolism for the growth of biomass:

$$HCO_3^- \xrightarrow{CA} OH^- + CO_2$$

The medium was changed every 5 days and pH of the old medium was recorded to calculate the amount of accumulated $OH^-$ ions.

| Day | pH | pOH | $[OH^-]$ or $[CO_2]$, µmol |
|---|---|---|---|
| 5 | 8.574 | 5.426 | 3.75 |
| 10 | 9.522 | 4.478 | 33.3 |
| 15 | 9.459 | 4.541 | 28.8 |
| 20 | 9.362 | 4.638 | 23.0 |
| 25 | 9.379 | 4.621 | 23.9 |
| 30 | 9.399 | 4.601 | 25.1 |

Every sample ($V_{sample}$ = 40 µL) was incubated in $V_{medium\ per\ sample}$ = 2 mL of BG11:ASNIII medium. Therefore, to calculate the amount of $CO_2$ sequestered by 1 ml of hydrogel:

$$n_{CO_2 sequestered} = c_{CO_2\ in\ medium} \times V_{medium\ per\ sample} \times \frac{1000\ \mu L}{40\ \mu L}$$

| Day | $CO_2$ sequestered with 1 mL of gel, µmol | Cumulative $CO_2$ sequestered with 1 mL of gel, µmol | Cumulative $CO_2$ sequestered with 1 mL of gel, mg |
|---|---|---|---|
| 5 | 0.19 | 0.19 | 0.0083 |
| 10 | 1.68 | 1.87 | 0.082 |
| 15 | 1.45 | 3.32 | 0.15 |
| 20 | 1.16 | 4.48 | 0.20 |
| 25 | 1.21 | 5.69 | 0.25 |
| 30 | 1.27 | 6.96 | 0.31 |

**Total equivalent $CO_2$ sequestration calculation in living materials**

To evaluate the mass of $CO_2$ converted into biomass and inorganic precipitates within hydrogel matrix we weighed the dry mass of randomly selected abiotic and biotic samples over the incubation period of 30 days. As the hydrogel is composed of 13.2 wt% Pluronic F127 and 7.3 wt% photo-cross-linkable F127-BUM, the non-cross-linked F127 diffused out over time and the dry mass of abiotic samples decreased to 45 ± 7% of the original mass. The same effect was expected for biotic samples. However, the final mass after 30 days of incubation was 81 ± 9% of the initial one as the reduction of polymer content was partially replaced by the generated biomass and inorganic precipitates.

$$mass_{abiotic} = mass_{F127} + mass_{F127-BUM}$$

$$mass_{biotic} = mass_{F127} + mass_{F127-BUM} + mass_{biomass} + mass_{precipitates}$$

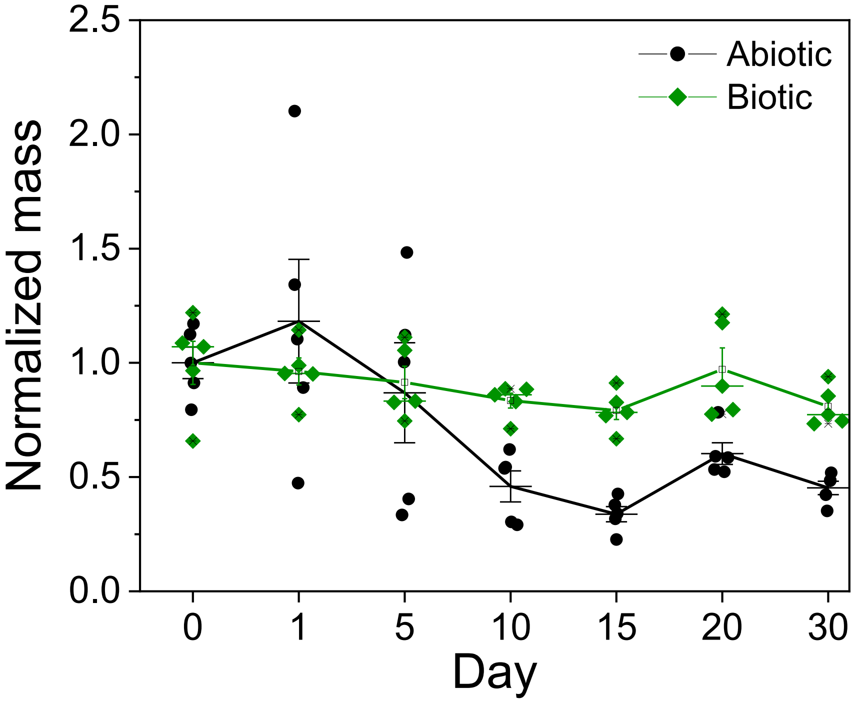

| $Mass_{abiotic}$, day 30, % | $Mass_{biotic}$, day 30, % | $\Delta(Mass_{biotic} - Mass_{abiotic})$, % |
|---|---|---|
| 45 ± 7 | 81 ± 9 | 36 |

To quantify the extent of insoluble precipitates, thermal decomposition was performed to remove the organic biomass and polymer matrix. Dry abiotic or biotic samples ($n_{samples}$=5 and $n_{replicates}$=5) with a total mass of 10-20 mg per replicate were placed in a crucible for thermal decomposition at 600 °C. The biotic sample mass after thermal decomposition, which corresponded to the mass of the insoluble carbonate precipitates, was 5 ± 2%. The remaining mass of the abiotic samples was 1 ± 1% and, therefore, we considered it insignificant.

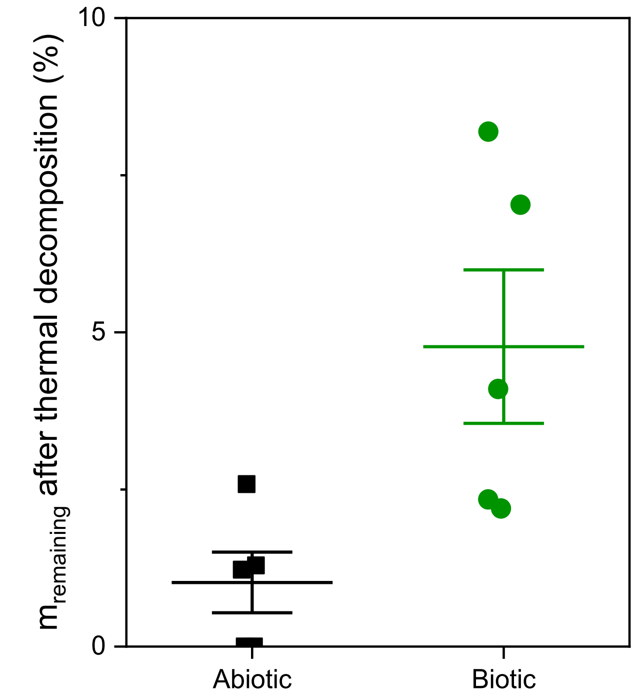

To convert this into the amount of sequestered $CO_2$ we used:

$V_{sample}$ = 40 μL – from the design of disc samples

$m_{dry\ sample}$ = 4 mg – from the measured average of biotic samples on day 30

$m_{inorganic\ precipitates\ per\ sample}$ = 0.2 mg – as 5% of sample mass based on the measured mass after thermal decomposition

number of samples per 1 mL of hydrogel = 1000 μL · $(0.04\ \mu L)^{-1}$ = 25

MW($CaCO_3$) = 100 g $mol^{-1}$

MW($MgCO_3$) = 84 g $mol^{-1}$

In case of a mixture of 70% $CaCO_3$ and 30% $MgCO_3$, as suggested by the XRD results in the calcite peak shift [4], average MW= 95.2 g $mol^{-1}$,

Then, the number of mole of $CaCO_3$ precipitates per 1 mL of hydrogel:

$$n\ (precipitates) = \frac{m_{inorganic\ precipitates\ per\ sample}}{\mathrm{M_w(CaCO_3)}} \cdot 25$$

| | Per sample, μmol | Per 1 mL of hydrogel, μmol |
|---|---|---|
| $CaCO_3$ | 2 | 50 |
| 70% $CaCO_3$ and 30% $MgCO_3$ | 2.1 | 52.5 |

This results in approximately 50 μmol of $CO_2$ sequestered as inorganic precipitates per 1 mL of living material or 2.2 ± 0.9 mg per 1 mL of living material.

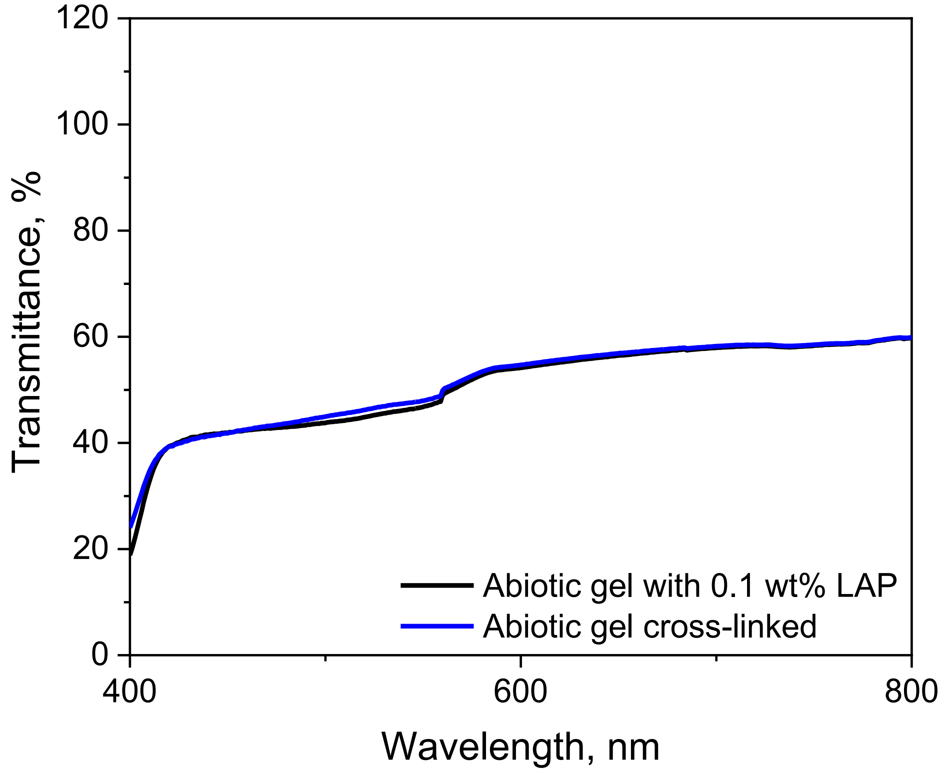

**Figure S1.** Transmittance of abiotic F127 based hydrogel with 0.1 wt% LAP photoinitiator. Cross-linking duration t = 2 min.

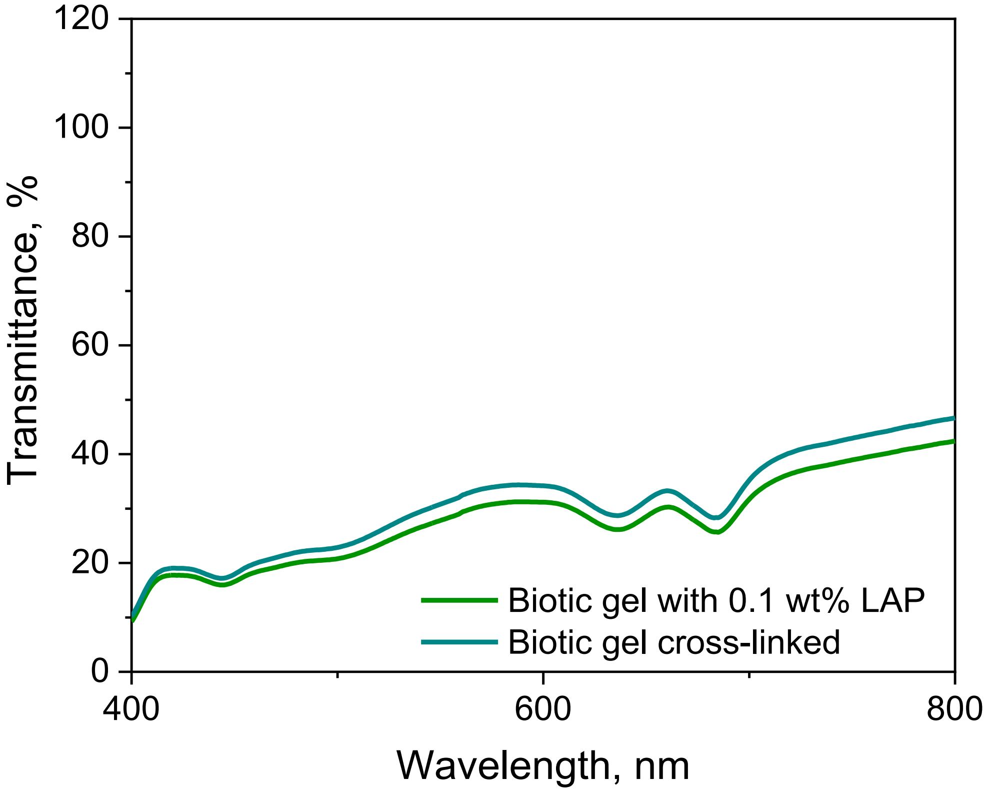

**Figure S2.** Transmittance of biotic F127 based hydrogel with 0.1 wt% LAP photoinitiator and PCC 7002 concentration equivalent to $OD_{730nm}$ = 0.8. Cross-linking duration t = 1 min.

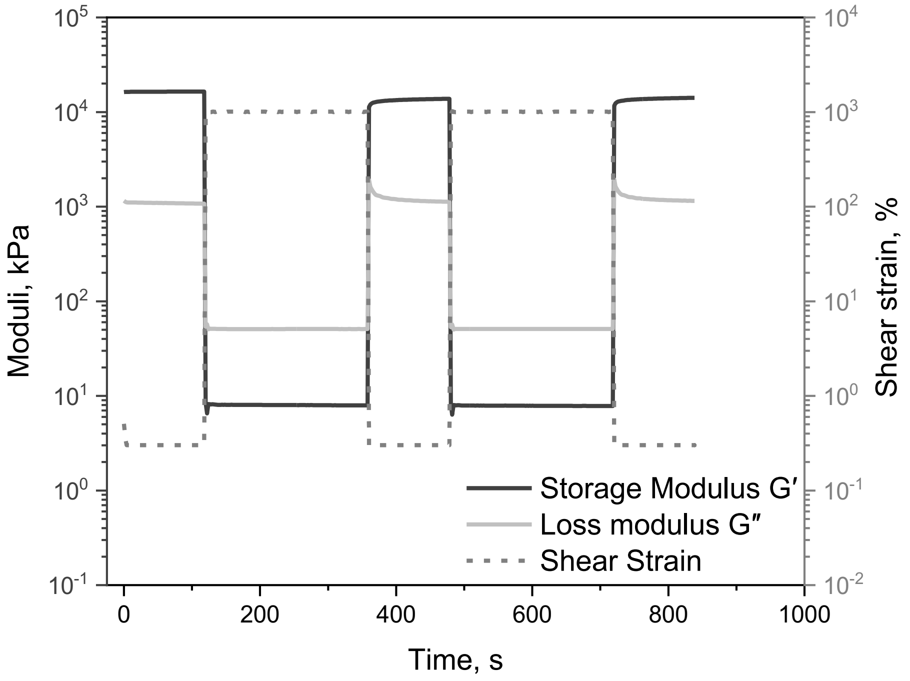

**Figure S3.** Step strain measurements for Pluronic F127 with alternating intervals at low (γ = 0.3%, ω = 10 rad s$^{-1}$) and high (γ = 1000%, ω = 10 rad s$^{-1}$) shear strain amplitude. F127 exhibited rapid and reproducible recovery of solid-like properties (G′ > G″; elastic recovery).

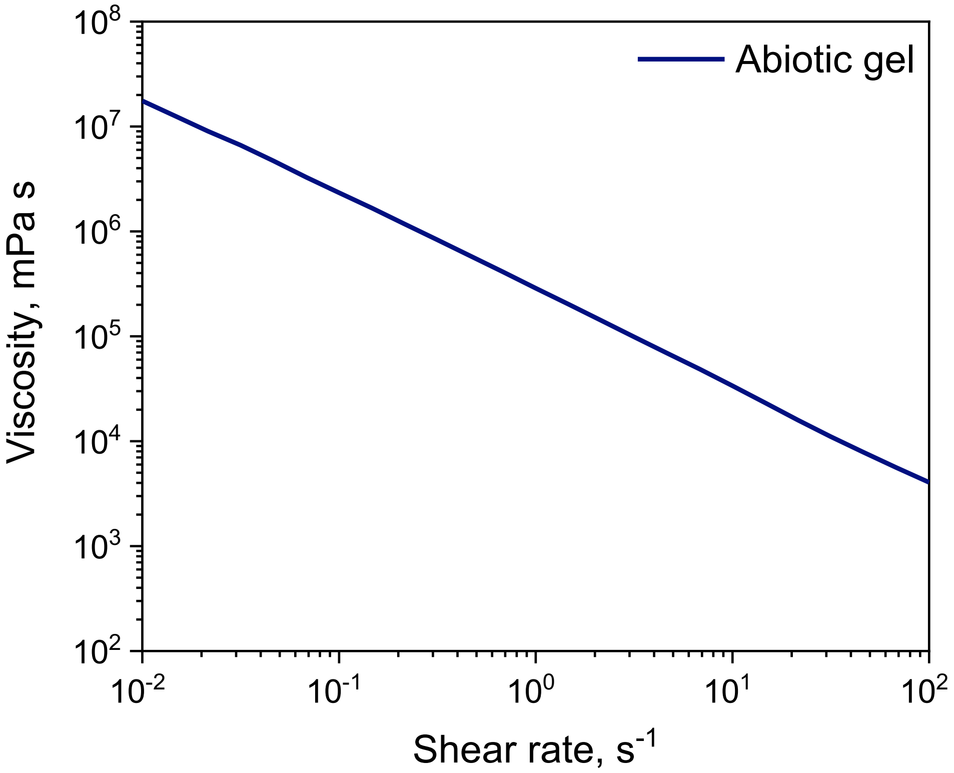

**Figure S4.** Shear rate ramp (dγ/dt = 0.1–100 $s^{-1}$) for Pluronic F127. F127 demonstrated a decrease in viscosity with increasing shear rate (shear-thinning).

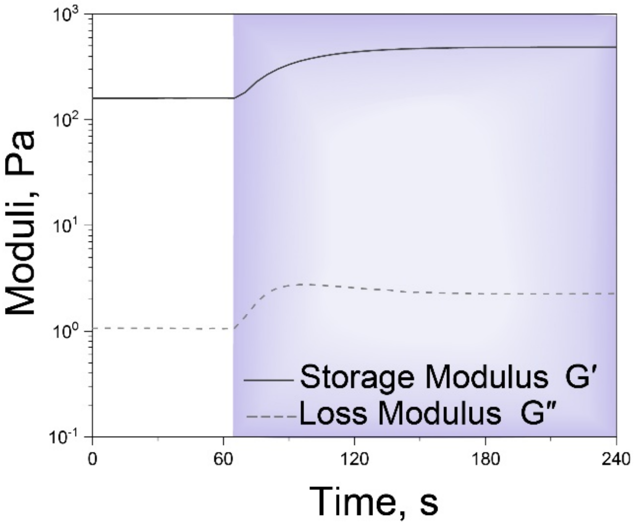

**Figure S5.** Photo-cross-linking of 13.2 wt% F127 and 7.3 wt% F127-BUM and 0.1 wt% LAP hydrogel using 405 nm (I = 8 mW $cm^{-2}$) light (light on in the region marked in violet).

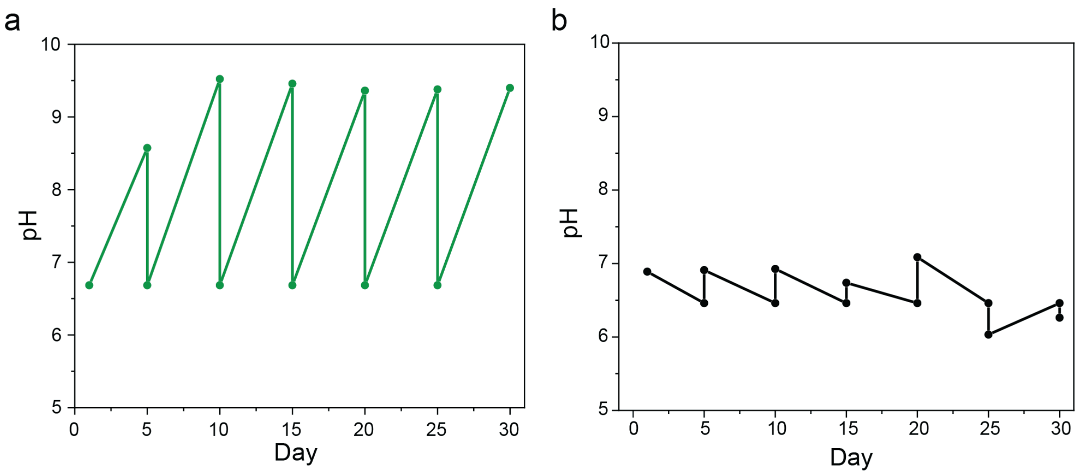

**Figure S6.** pH change of the growth medium for a) biotic and b) abiotic samples throughout the incubation period of 30 days. The pH value of the medium returned to 6.5 every 5 days due to the medium change.

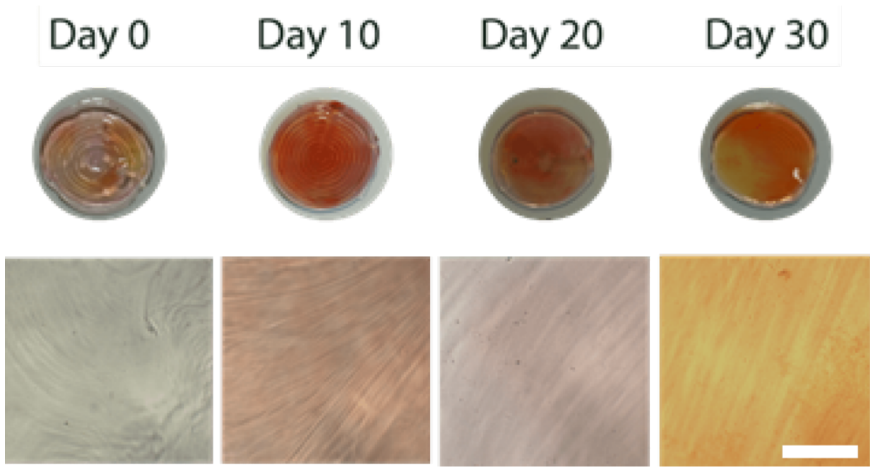

**Figure S7.** Optical and microscopic images of Alizarin red S staining on abiotic samples. Scale bar, 100μm.

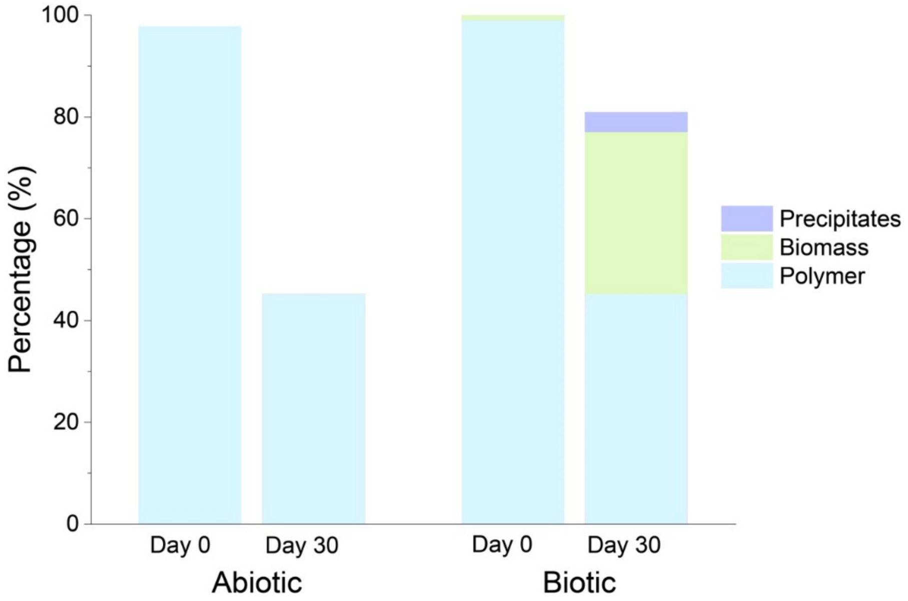

**Figure S8.** Composition of abiotic and biotic samples on day 0 and day 30. The dry mass of both abiotic and biotic day 30 samples was normalized to the mass on day 0. The decrease in mass in the abiotic samples was due to the diffusion of unfunctionalized Pluronic F127 into the cell culture medium.

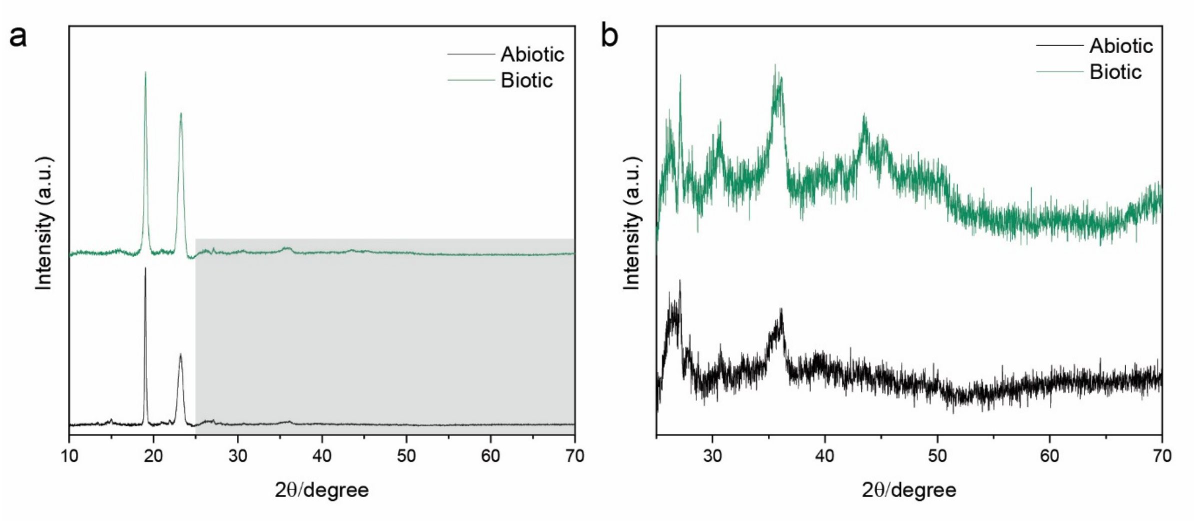

**Figure S9.** XRD analysis of the abiotic and biotic samples after 30 days of incubation. The samples were first lyophilized to remove any water content, followed by homogenizing in 1 mL of absolute ethanol for 2 minutes. The major peaks at 18 and 23 correspond to the Pluronic F127 micelles and the calcite peaks are not prominent due to the strong signal from the F127 micelles. (a) XRD spectra over 2θ of 10 - 70°, (b) zoomed-in XRD spectra between a 2θ of 25 and 70°. The signal from the F127-based matrix was much stronger than the one of the precipitates and the signal at 31° could be associated with the calcite peak post thermal decomposition.

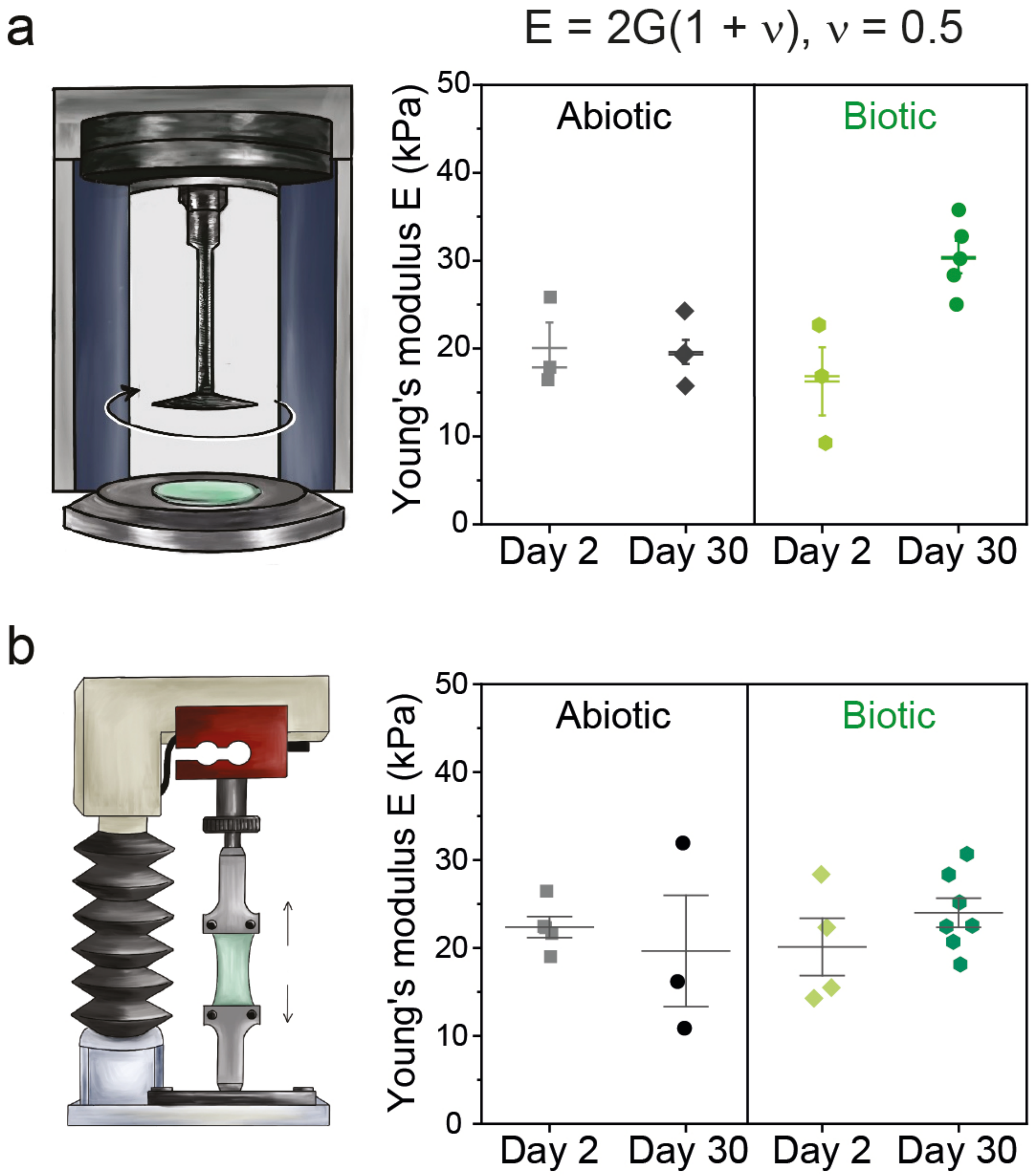

**Figure S10.** Hydrogel Young's modulus: (a) calculated based on shear modulus obtained with shear rheometer by assuming a Poisson's ratio of $\nu = 0.5$ and (b) by tensile test. Both the tensile test and rheology showed similar Young's modulus values for the biotic and abiotic samples on day 2 and day 30.

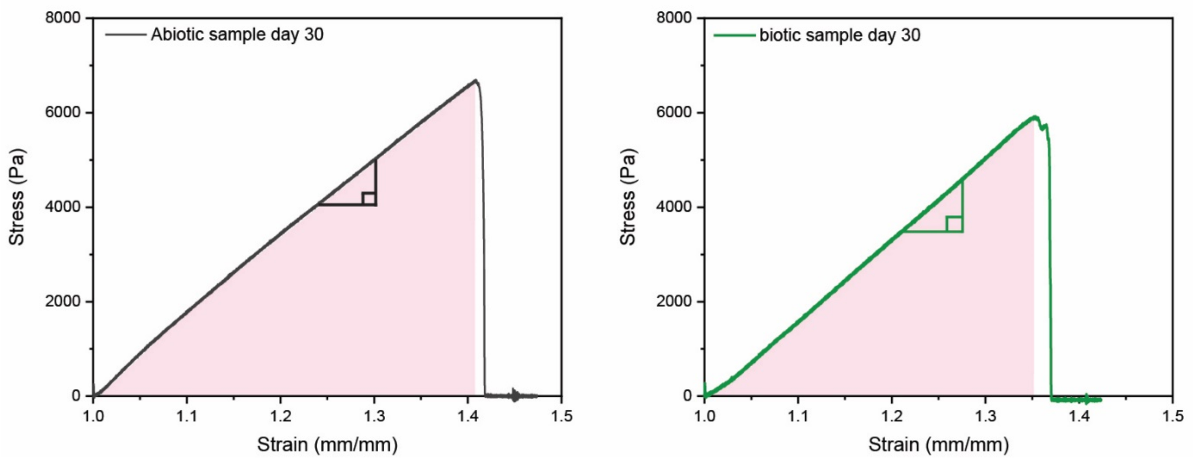

**Figure 11.** Stress-strain curve of day 30 abiotic and biotic samples obtained from uniaxial tensile test. The slope of the stress-strain curve was calculated as the Young's modulus of the samples and the area under the curve was integrated as the material's toughness.

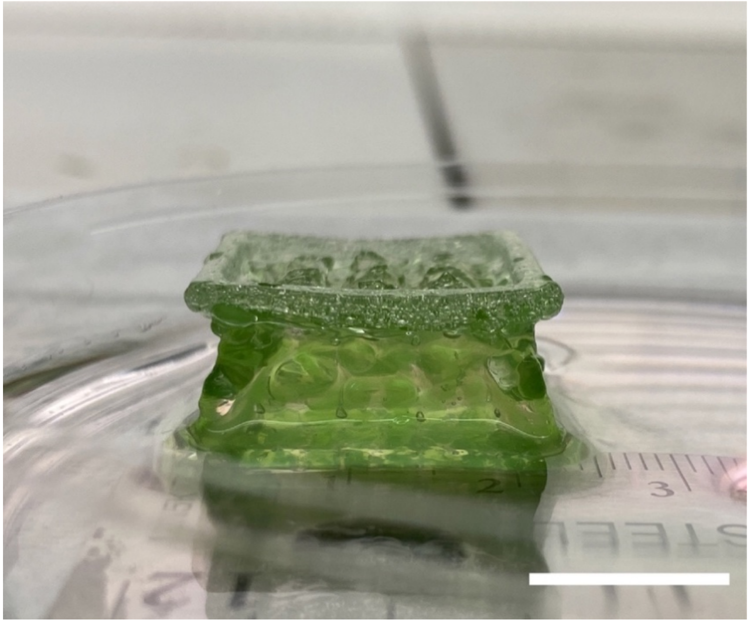

**Figure S12.** Designed BCC structure that actively transports medium upwards**.** Scale bar, 1 cm.

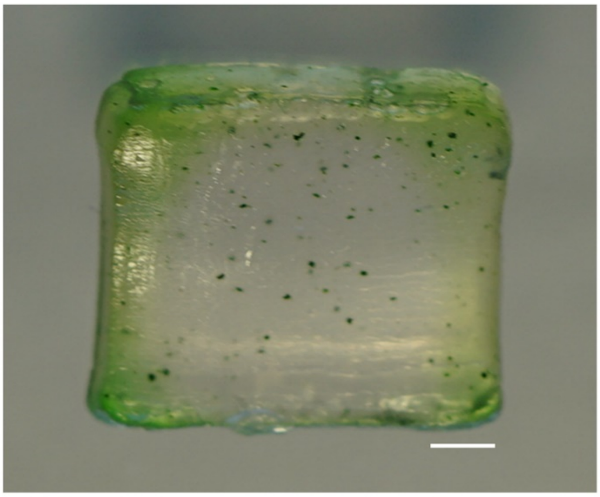

**Figure S13.** The maximum thickness of the living material to maintain bacteria viability was found to be approximately 2.5 mm from each side (indicated by the green color) and thus a total depth of 5 mm from both sides. Scale bar, 2 mm.

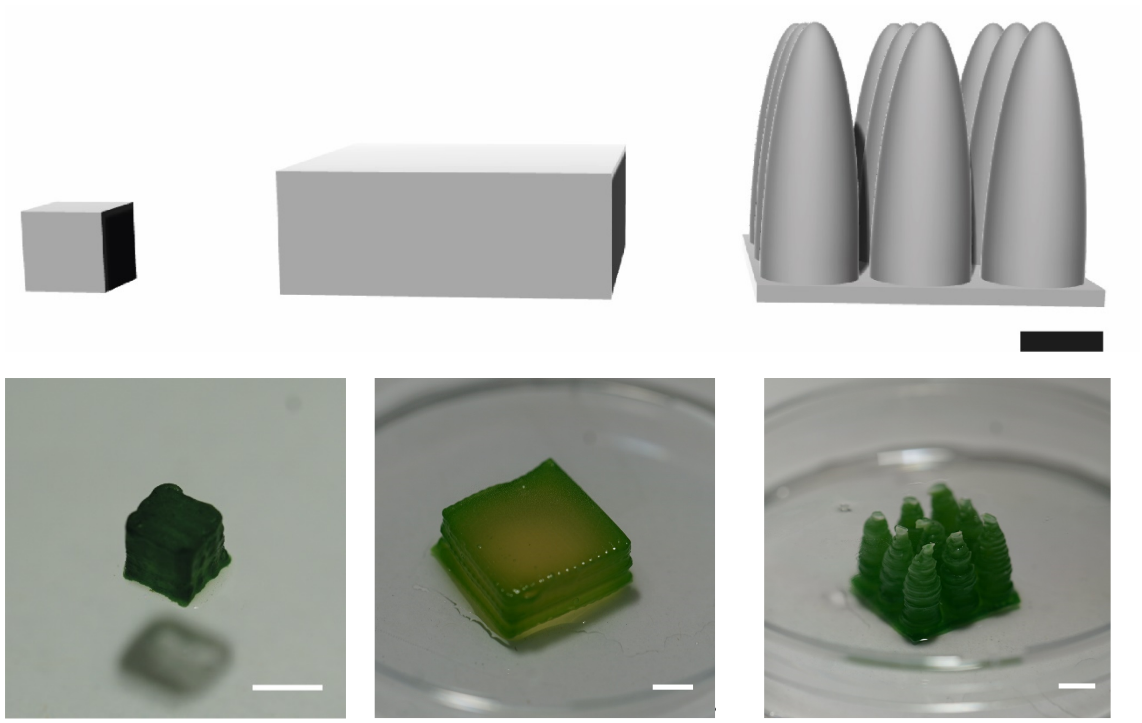

5 x 5 x 5 mm

$0.5\ g_{material}\ cm^{-2}$

~100% viable volume

20 x 20 x 7.5 mm block

$0.75\ g_{material}\ cm^{-2}$

<<100% viable volume

20 x 20 mm Pillar array

$0.75\ g_{material}\ cm^{-2}$

~100% viable volume

**Figure S14.** 3D models of bulk (left and middle) and pillar array structure (right) and corresponding printed structures. Printed 5 x 5 x 5 mm bulk volume had ~ 100 % viable volume on day 16. However, the 20 x 20 x 7.5 block was not fully viable on day 5 and, therefore, pillar array design was employed to increase the viable volume of material per area. The pillar array designed with the same material volume and surface coverage had ~ 100 % viability on day 5. Scale bars, 0.5 cm.

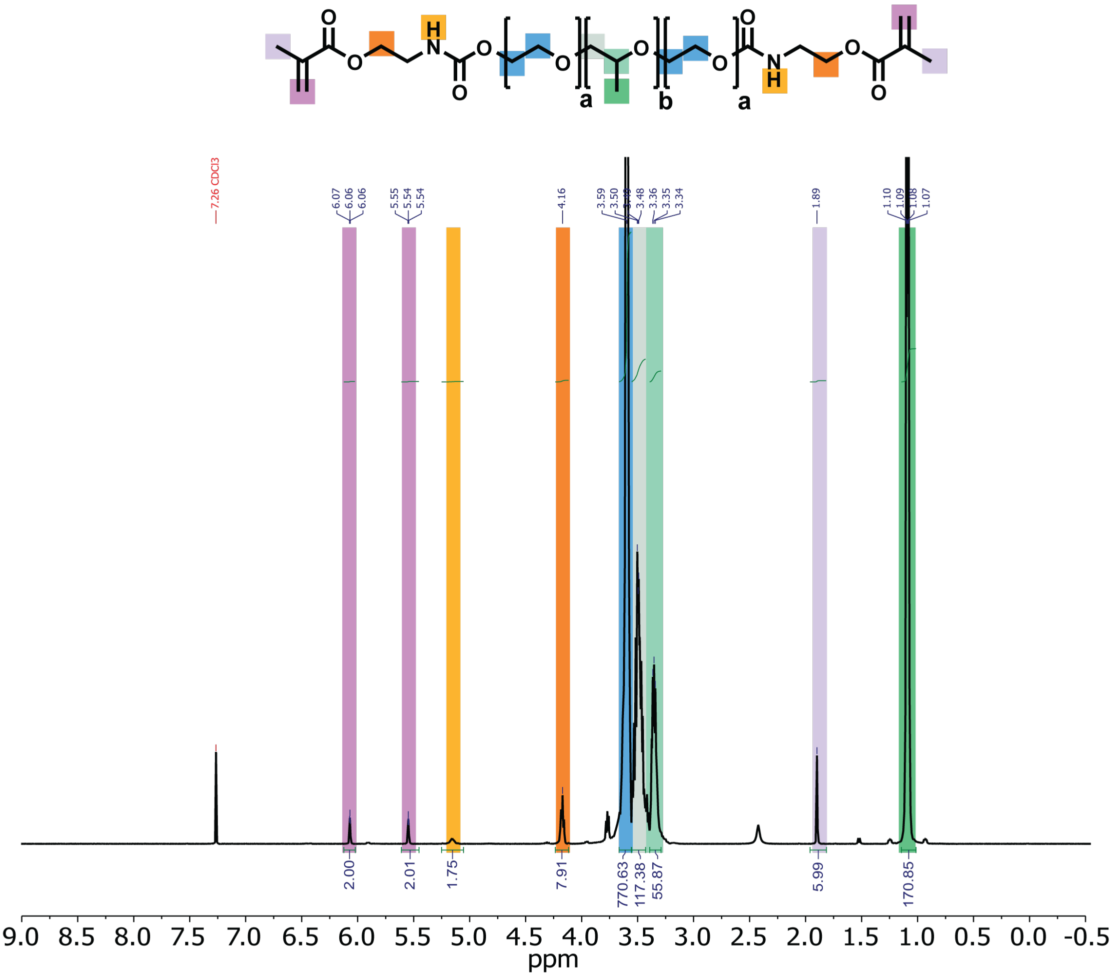

**Figure S15.** $^{1}$H-NMR spectrum of F127-BUM in $CDCl_3$.

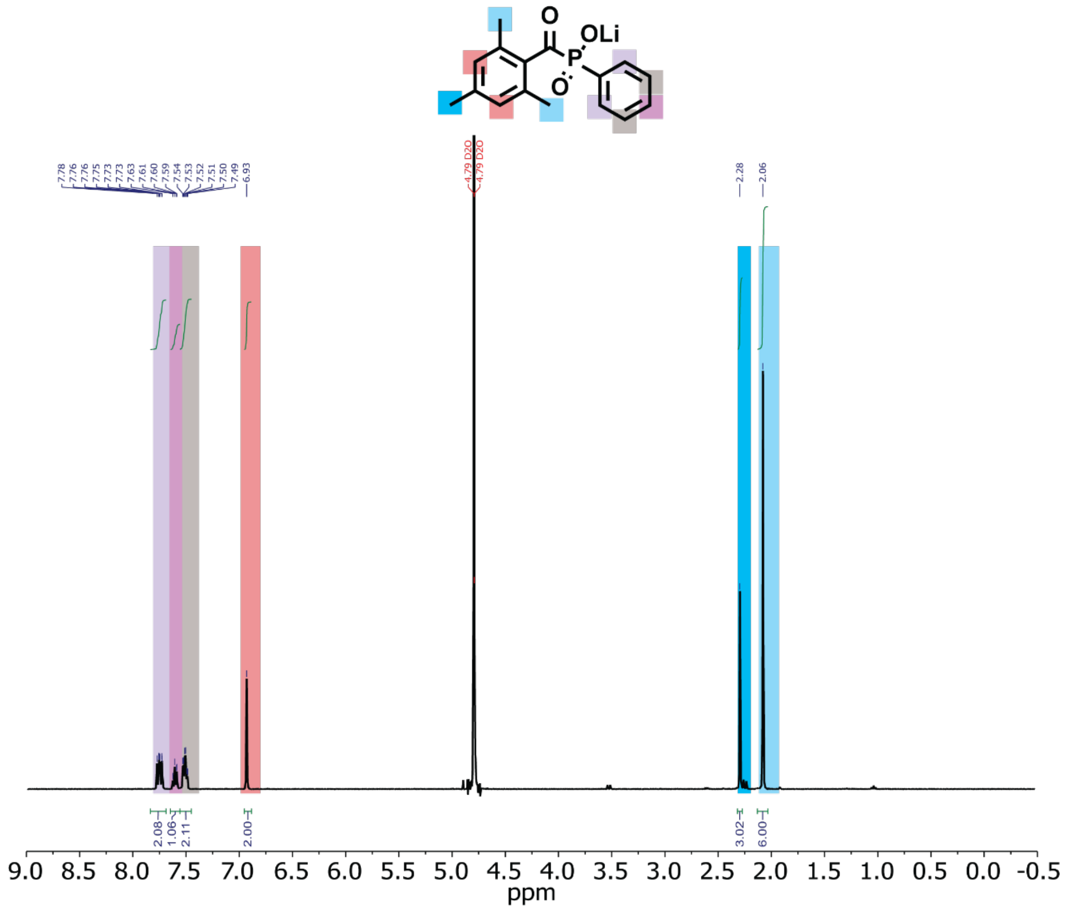

**Figure S16.** 1H-NMR spectrum of LAP in $D_2O$.

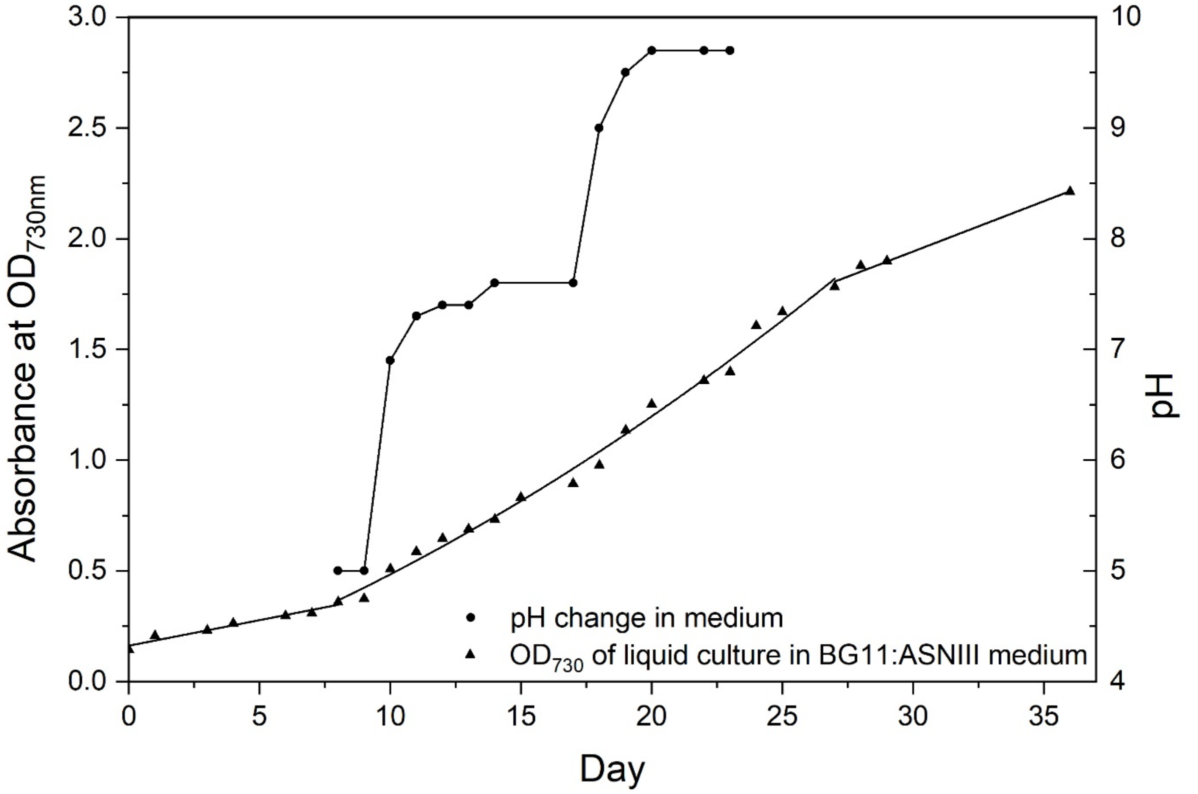

**Figure S17.** Growth curve of PCC 7002 in BG11: ASNIII medium and pH change of the liquid culture.

**Table S1. ASNIII medium components**

| Component | Concentration [g $L^{-1}$] |
|---|---|
| NaCl | 25.0 |
| $MgSO_4$ anhydrous | 1.78 |
| KCl | 0.50 |
| $NaNO_3$ | 2.25 |
| $MgCl_2 \cdot 6H_2O$ | 2.00 |
| $K_2HPO_4$ anhydrous | 0.043 |
| $CaCl_2$ anhydrous | 0.405 |
| Citric Acid | 0.009 |
| Ferric ammonium citrate | 0.009 |
| EDTA (disodium magnesium) | 0.001 |
| Milli-Q water | to 1 L |

**Table S2. Shear-thinning parameters**

| Sample | K, Pa $s^{-n}$ | n | $r^2$ |
|---|---|---|---|
| Abiotic hydrogel | 287±7 | 0.091±0.005 | 0.99 |

**Table S3. Compressive modulus of photosynthetic living materials after 400 days of incubation**

| s/n | Sample thickness [mm] | Compressive modulus [kPa] |
|---|---|---|
| 1 | 3.871 | 120.9 |
| 2 | 3.966 | 105.1 |
| 3 | 3.218 | 102.8 |
| 4 | 3.727 | 112.8 |
| 5 | 3.765 | 113.7 |

**Table S4. $CO_2$ sequestration per gram of hydrogel material after 400 days of incubation, the mass of $CaCO_3$ precipitates were obtained after thermal decomposition at 600 °C**

| s/n | Sample wet mass [mg] | $CaCO_3$ mass [mg] | $CO_2$ sequestration per gram of hydrogel material [mg · $g^{-1}$] |
|---|---|---|---|
| 1 | 206 | 339.9 | 34.0 |
| 2 | 60.4 | 215.5 | 21.6 |
| 3 | 64.7 | 258.3 | 25.8 |
| 4 | 114.5 | 355.9 | 30.4 |
